# Estimation of the probability of epidemic fade-out from multiple outbreak data

**DOI:** 10.1101/2021.06.01.446666

**Authors:** Punya Alahakoon, James M. McCaw, Peter G. Taylor

## Abstract

Deterministic epidemic models, such as the *SIRS* model or an *SIR* model with demography, that allow for replenishment of susceptibles typically display damped oscillatory behaviour. If the population is initially fully susceptible, once an epidemic takes off a distinct trough will exist between the first and second waves of infection where the number of infectious individuals falls to a low level. Epidemic dynamics are, however, influenced by stochastic effects, particularly when the number of infectives is low. At the beginning of an epidemic, stochastic die-out is possible and well characterised through use of a branching process approximation to the full non-linear stochastic dynamics. Conditional on an epidemic taking off, stochastic extinction is highly unlikely during the first epidemic wave, but the probability of extinction increases again as the wave declines. Extinction during this period, prior to a potential second wave of infection, is defined as ‘epidemic fade-out’. We consider a set of observed epidemics, each distinct and having evolved independently, in which some display fade-out and some do not. While fade-out is necessarily a stochastic phenomenon, in general the probability of fade-out will depend on the model parameters associated with each epidemic. Accordingly, we ask whether time-series data for the epidemics contain sufficient information to identify the key driver(s) of different outcomes—fade-out or otherwise—across the sub-populations supporting each epidemic. We apply a Bayesian hierarchical modelling framework to synthetic data from an *SIRS* model of epidemic dynamics and demonstrate that we can 1) identify when the sub-population specific model parameters supporting each epidemic have significant variability and 2) estimate the probability of epidemic fade-out for each sub-population. We demonstrate that a hierarchical analysis can provide more accurate and precise estimates of the probability of fade-out than is possible if considering each epidemic in isolation. Our methods may be applied more generally, to both epidemiological and other biological data to identify where differences in outcome—fade-out or recurrent infection/waves are purely due to chance or driven by underlying changes in the parameters driving the dynamics.

## 1 Introduction

Recurrent infectious diseases such as measles and chickenpox in populations where susceptibles replenish have been of mathematical and epidemiological interest for many years (Bartlett, 1957, 1960; Bjørnstad, Finkenstädt, & Grenfell, 2002; Keeling & Grenfell, 1997; London & Yorke, 1973). Replenishment of susceptibles can happen through births, immigration, or loss of immunity to the disease. Following the introduction of the pathogen to a fully susceptible population, when modelled deterministically, prevalence of infection can exhibit damped oscillatory behaviour. During the troughs preceded by the outbreaks of infection, prevalence drops to a low level until the infection takes off again (with an effective reproduction number, *R*_eff_ *>* 1) to generate the next outbreak.

Disease dynamics are however, impacted by stochasticity, particularly at low levels of prevalence. This gives rise to variability in the epidemic trajectory as well as the possibility for extinction of the pathogen from the population (Lloyd, 2004). At the beginning of the epidemic when the prevalence is low, the pathogen can “fade-out” without infecting a significant number of people. This type of initial extinction is well studied using branching processes (Allen, 2015; Allen & van den Driessche, 2013; Britton et al., 2019).

If the initial fade-out does not occur, the epidemic will take off and extinction during the outbreak is highly unlikely. As the epidemic advances in time and the number of susceptibles depletes, the effective reproduction number reduces. Once it drops below one, the epidemic is not self sustaining and prevalence begins to decline, ultimately reaching a low level(Lloyd, 2004). Once prevalence is at a low level, stochastic effects become apparent and the probability of extinction begins to increase (Anderson & May, 1979). This probability is likely to reach a maximum during the time period in which, under a deterministic analysis, the first trough between outbreaks would have occurred (Lloyd, 2004). If extinction does not occur, the replenishment of susceptibles will increase the effective reproduction number (*R*_eff_ > 1), and a second outbreak may be initiated.

When stochastic effects drive the prevalence of infection to zero before the potential second outbreak can take place, it is known as an epidemic fade-out. The notion of epidemic fade-out can be seen in studies by Anderson and May (1979, 1992); Ballard, Bean, and Ross (2016); Bartlett (1957, 1960); Camacho et al. (2011); Camacho and Cazelles (2013); Keeling and Grenfell (1997); Lloyd-Smith et al. (2005); Meerson and Sasorov (2009); van Herwaarden (1997).

We consider multiple outbreaks of a recurrent infectious disease that take place independently in a set of sub-populations. The sub-populations we consider are in contained environments and separated from each other. Examples of such sub-populations are islands, ships, or individuals in a clinical trial. There is no movement between these sub-populations and transmission of the disease in one sub-population does not influence the other. Different transmission patterns are likely to arise from stochastic effects and variations in the sub-populations’ disease-related factors such as transmission rates, infectious and/or latent periods or disease resistance, and other factors such as social, cultural, demographic, and geographic factors. Infectious disease model parameters of these sub-populations will therefore be rarely identical. Identifying and understanding the consequences of the existence of this variability is the major goal of this study.

In this study, we investigate if it is possible to identify whether differences in observing epidemic fade-outs in multiple sub-populations can be attributed just to stochastic effects or if they are influenced by different characteristics of each sub-population. We apply a simulation-based approach by generating synthetic data for two populations from a stochastic *SIRS* model (introduced in Section 2). The data for one population is generated by independent stochastic *SIRS* models with identical parameters across the sub-populations, while the data for the second population is generated using stochastic *SIRS* models with different parameters across the sub-populations. The sub-populations may or may not display an epidemic fade-out. We apply a stochastic Bayesian hierarchical framework to estimate model parameters, introducing an efficient two-step Approximate Bayesian Computation (ABC) methodology. Using the estimated model parameters within the hierarchical framework, we illustrate that we are able to identify, when present, the level of variability between the model parameters of the sub-populations. After estimating the parameters, we characterise their influence on the probability of epidemic fade-out.

## 2 Background

### 2.1 The Markovian *SIRS* model in a closed sub-population

For a well-mixed sub-population of size *N, S*(*t*), *I*(*t*) and *R*(*t*) denote the numbers of susceptible, infectious, and recovered individuals at time *t*. The model is parameterised by *β*, the contact rate, *γ*, the rate of recovery, and *µ*, the waning immunity rate (Figure 1). Noting that *R*(*t*) = *N— S*(*t*) *−I*(*t*), a stochastic SIRS system can be modelled with a continuous-time Markov chain with bi-variate states (*S*(*t*), *I*(*t*)) and transition rates presented in Table 1.

**Table 1:**
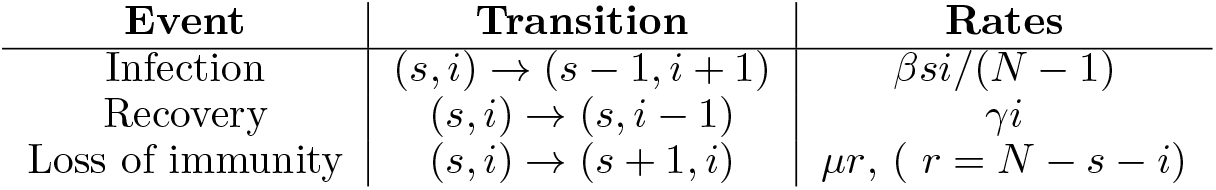
Transition rates of an *SIRS* model

| Event | Transition | Rates |
| --- | --- | --- |
| Infection | $(s, i) \rightarrow (s - 1, i + 1)$ | $\beta si / (N - 1)$ |
| Recovery | $(s, i) \rightarrow (s, i - 1)$ | $\gamma i$ |
| Loss of immunity | $(s, i) \rightarrow (s + 1, i)$ | $\mu r, (r = N - s - i)$ |

**Figure 1:**
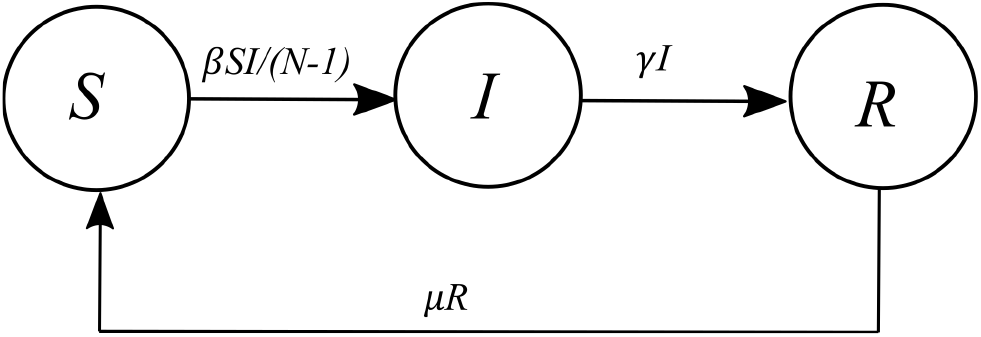
Diagram of an *SIRS* model structure.

### 2.2 Structure of the study

We consider the following two scenarios regarding the variability between model parameters of the sub-populations in which an epidemic occurs.

Scenario 1: The sub-populations all have identical parameters.

Scenario 2: There exists significant variability between the sub-population specific parameters.

In any given sub-population, the epidemic may fade-out or not. If all the sub-populations had identical model parameters (under Scenario 1 above), we can conclude that epidemic fade-outs (or non-fade-outs) in the sub-populations occur due to stochastic fluctuations only. However, if the sub-populations fall into Scenario 2, we cannot make such direct conclusions as variations in the occurrence of epidemic-fade-out could be a result of the differences in parameters as well as stochastic effects.

In Section 3, we will first discuss how parameter estimation for stochastic models can be carried out using hierarchical Bayesian estimation techniques. Then we will construct two synthetic datasets reflecting the two scenarios above and carry out parameter estimation and explore how we can effectively categorise the populations based on the level of variability in the model parameters.

The second part of this study is devoted to identifying the estimated parametric effects on the probability of epidemic fade-out. We explore how to incorporate the information obtained from the inference to draw informed conclusions about the probability of epidemic fade-out in the two populations.

Using an additional synthetic dataset, we will compare the estimates obtained with and without a hierarchical modelling framework and illustrate that making use of a hierarchical modelling framework yields accurate results as a reference to the true parameters and probability of epidemic fade-out.

## 3 Materials and Methods

### 3.1 A Bayesian hierarchical modelling approach

For *k* = 1, …, *K* sub-populations, let ***X***_*k*_(*t*) = ((*S*_*k*_(*t*), *I*_*k*_(*t*))) be a continuous-time Markov chain with transition rates given in Table 1 and ***X*(*t*)** = (***X***_**1**_**(*t*)**, …, ***X***_***K***_**(*t*)**). We consider a hierarchical modelling approach to estimate the parameters of the *K* sub-populations where each sub-population has the same *SIRS* model structure with model parameters drawn from a common distribution. Some examples on the usage of hierarchical models and their applications are Gelman, Hill, and Yajima (2012); Kwok and Lewis (2011).

We will consider a hierarchical Bayesian modelling framework with three levels. Level I represents prevalence data at each sub-population. For sub-population *k* with parameters ***θ***_*k*_ (*k* = 1, 2, …, *K*), our data consists of the vector ***y***_***k***_ = (*I*_*k*_(1), *I*_*k*_(2) …, *I*_*k*_(*T*_*k*_)) at *T*_*k*_ discrete time points and ***y*** = (***y***_**1**_, ***y***_**2**_, …, ***y***_***K***_). Level II represents the structural relationship between the sub-populations in terms of a conditional prior, *p*(***θ***_*k*_| ***ψ***), that further depend on the hyper-parameters ***ψ***. In level II, we assume that the sup-population specific parameters are a random sample from the conditional prior distribution. Prior distributions of the hyper-parameters, that is, hyper-prior distributions *p*(***ψ***), are presented in Level III (Gelman et al., 2013).

The joint posterior distribution for a population consisting of *K* sub-populations is,

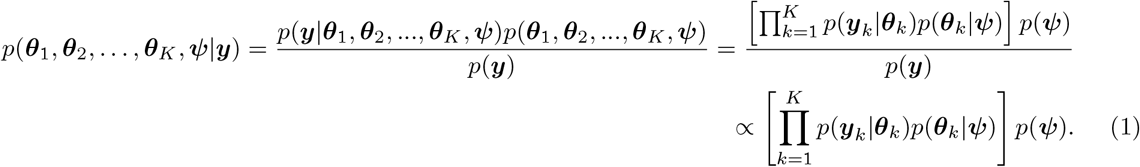

Estimating this posterior is often difficult in the sense that for most models, including epidemic models, a closed-form expression for the posterior distribution does not exist. Therefore, we resort to methods that enable us to sample from the marginal posterior. Since the likelihood of these stochastic epidemic models is also intractable, we further resort to Approximate Bayesian Computation (ABC) methods that avoid calculating the likelihood when sampling from the posterior (Sisson, Fan, & Beaumont, 2018).

### 3.2 Parameter estimation using ABC methods

In this subsection, we consider applying a basic ABC algorithm for the *k*th sub-population independently and in the next subsection, we examine how to extend these methods to all the *K* sub-populations within a hierarchical setting using conventional methods as well as using a novel computationally efficient method.

The simplest form of an ABC algorithm to obtain a sample from the posterior *p*(***y***_*k*_ |***θ***_*k*_) is as follows: first, sample a parameter set ***θ***_***k***_^(*j*)^ (*j* = 1, 2, … *N*) from the prior distribution *p*(***θ***_*k*_). Next, generate a dataset 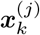 from the model described by 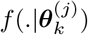. Third, use 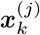 to construct generated summary statistics 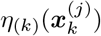 and use ***y***_*k*_ to construct observed summary statistics *η*_(*k*)_(***y***_*k*_). Finally, using a distance *ρ*_(*k*)_(.,.), retain 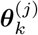 if 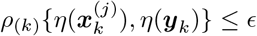, for some small tolerance value *∊ >* 0 (Marin, Pudlo, Robert, & Ryder, 2012).

We use the observed number of infectious individuals recorded at discrete times as the observed summary statistics (examples: McKinley, Cook, and Deardon (2009); Minter and Retkute (2019)). We use the Doob-Gillespie algorithm (Doob, 1945; Gillespie, 1977) to generate data. We take the number of infectious individuals retained at discrete times 1, 2, …, *T*_*k*_ from a generated sample for the *k*th sub-population as the generated summary statistic *η*_(*k*)_(***x***_*k*_) = ***x***_*k*_ = (*x*_*k*_(1), *x*_*k*_(2), …, *x*_*k*_(*T*_*k*_)). We further take the square root of the Euclidean distance,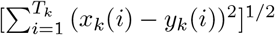, as the distance metric, *ρ*_(*k*)_(***x***_*k*_, ***y***), for the *k*th sub-population.

### 3.3 A novel method to estimate parameters of a hierarchical model

We first focus on applying a hierarchical modelling framework within a conventional ABC algorithm. First, a vector of hyper-parameters, ***ψ***^**′**^ are sampled from the hyper-prior distribution, *p*(***ψ***). Next, vectors of sub-population specific parameters, ***θ***^**′**^_*k*_, are sampled from the conditional prior *p*(***θ***_*k*_ | ***ψ***^**′**^). The ABC acceptance-rejection criteria must be applied to all *K* vectors of sub-population specific parameters in unison to either accept or reject ***ψ***^**′**^ and ***θ***^**′**^_*k*_. Each iteration would be computationally expensive and inefficient in the sense that as the number of sub-populations become larger, the whole set of parameters 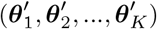 would be rejected if any one 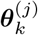 does not fulfil the chosen ABC criteria. A similar observation had been made by Bazin, Dawson, and Beaumont (2010).

Therefore, we propose an efficient two-step method to estimate the parameters of a hierarchical model. While addressing the drawbacks of the previous methods, this method is also easy to computationally implement and the overall performance can be accelerated by parallel computing techniques.

The first step of this method consists of estimating the hyper-parameters. The second step estimates the sub-population specific parameters given the estimated hyper-parameters. This approach is similar to the two-step algorithm introduced by Bazin et al. (2010). Bazin et al. (2010) used symmetric summary statistics (a function of all sub-populations together) to infer the hyper-parameters in the first step and unit-specific summary statistics (for a sub-population) to infer sub-population specific parameters. This method requires finding two types of distinct summary statistics —at the hyper parametric and sub-population specific parameter level. This stringent requisite can be problematic when observed data are already in a lower dimension (such as the prevalent data we study) or when the summary statistics that are either sufficient or approximately sufficient are very limited. To address this issue, we introduce a two-step method that make use of sub-population specific summary statistics, *η*_(*k*)_(.), in both steps. The two steps of our methodology are,

#### Step 1: Estimating hyper-parameters

a. Choose a “reasonable prior” for the distributions of the sub-population specific parameters ***θ***_*k*_. For each sub-population *k*, use this prior and an ABC-based algorithm independently to get *N*_1_ samples from the marginal posterior of ***θ***_*k*_ (see Supplementary materiel S1 for further details).
b. With an approximate estimator for the likelihood at the hyper-parametric level (i.,e,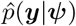) use the *KN*_1_ posterior samples generated in Step 1(a) to replace the likelihood function of an MCMC (Metropolis, Rosenbluth, Rosenbluth, Teller, & Teller, 1953) algorithm to obtain a sample of size *N*_2_ from the posteriors of the hyper-parameters, ***ψ*** (use the procedure described in Supplementary Material S1.1).

#### Step 2: Estimating sub-population specific parameters

a. Use the basic ABC algorithm to obtain samples of size *KN*_2_ from the sub-population specific posteriors, ***θ***_*k*_ given the hyper-parameters. In this step, sampling must be done from the conditional prior, *p*(***θ***_*k*_|***ψ***).

This method is efficient. In Step 1 (a) the sub-population-wise ABC-based algorithms can be run in parallel. Some examples of ABC-based algorithms in the literature include the algorithms by Beaumont, Cornuet, Marin, and Robert (2009); Sisson, Fan, and Tanaka (2007); Toni, Welch, Strelkowa, Ipsen, and Stumpf (2009). An ABC algorithm itself is inherently parallelizable and therefore, further computational efficiency can be achieved when sub-populations are treated independently at this step. There are also software packages readily available that can easily undertake this step. Part (b) of Step 1 is similar to particle-marginal methods (Andrieu, Doucet, & Holenstein, 2010) where the likelihood is replaced with a Monte Carlo estimate. Step 2 begins with samples from the marginal posteriors of the hyper-parameters. Again, given the hyper-parameters, samples from the posteriors of the sub-populations can be obtained by using a basic ABC algorithm independently (and hence can be done in parallel). Theory and the related algorithms of the two-step method are explained in detail in Supplementary Material S1.

### 3.4 Synthetic data generation

We constructed data for two distinct populations, A and B. Each consisted of 15 sub-populations of 1000 individuals. We simulated distinct *SIRS* epidemic dynamics using parameters selected as we describe below. For all the sub-populations, we consider observations of prevalence made at regularly spaced discrete time intervals over a 30-day time-period without any observational noise.

We generated synthetic data for all the sub-populations of Population A by simulating the stochastic *SIRS* model with identical parameters (*β*_*k*_, *γ, µ*) = (2, 1, 0.06) (for *k* = 1, 2, …, 15), starting with one infectious individual. We generated sample paths for all the sub-population up-to 30 days and we retained the number of infectious individuals on each day. If the number of infectious individuals of the sample path became zero before an initial outbreak took place, the sample path was removed and new sample paths were repeatedly generated until an initial outbreak was present (see Supplementary Material S3.1 to see the criterion we used to decide if an initial outbreak was present in the synthetic data).

In generating a synthetic dataset for Population B, we kept the values of *γ* and *µ* constant at 1 and 0.6 respectively, but we sampled the values of *β*_*k*_ (for *k* = 1, 2, …, 15) independently from a normal distribution with mean 2 and standard deviation 0.5, truncated to the interval (1, 10). The true mean and the standard deviation of this truncated normal distribution are 2.0276 and 0.4708 respectively. Given the model parameters for each sub-population, we carried out a synthetic data generation process similar to that of Population A.

Figures 2 and 3 show the synthetic datasets for the two populations. Table 2 shows the summary statistics of the sub-population specific parameters from which the data were generated in Population B.

**Table 2:** Summary statistics of the *β*_*k*_ (for *k* = 1, 2, …, 15) values of the 15 sub-populations in Population B.

| Mean | Median | Standard deviation | Minimum | Maximum |
| --- | --- | --- | --- | --- |
| 2.1805 | 2.2423 | 0.4821 | 1.2126 | 2.8778 |

**Figure 2:**
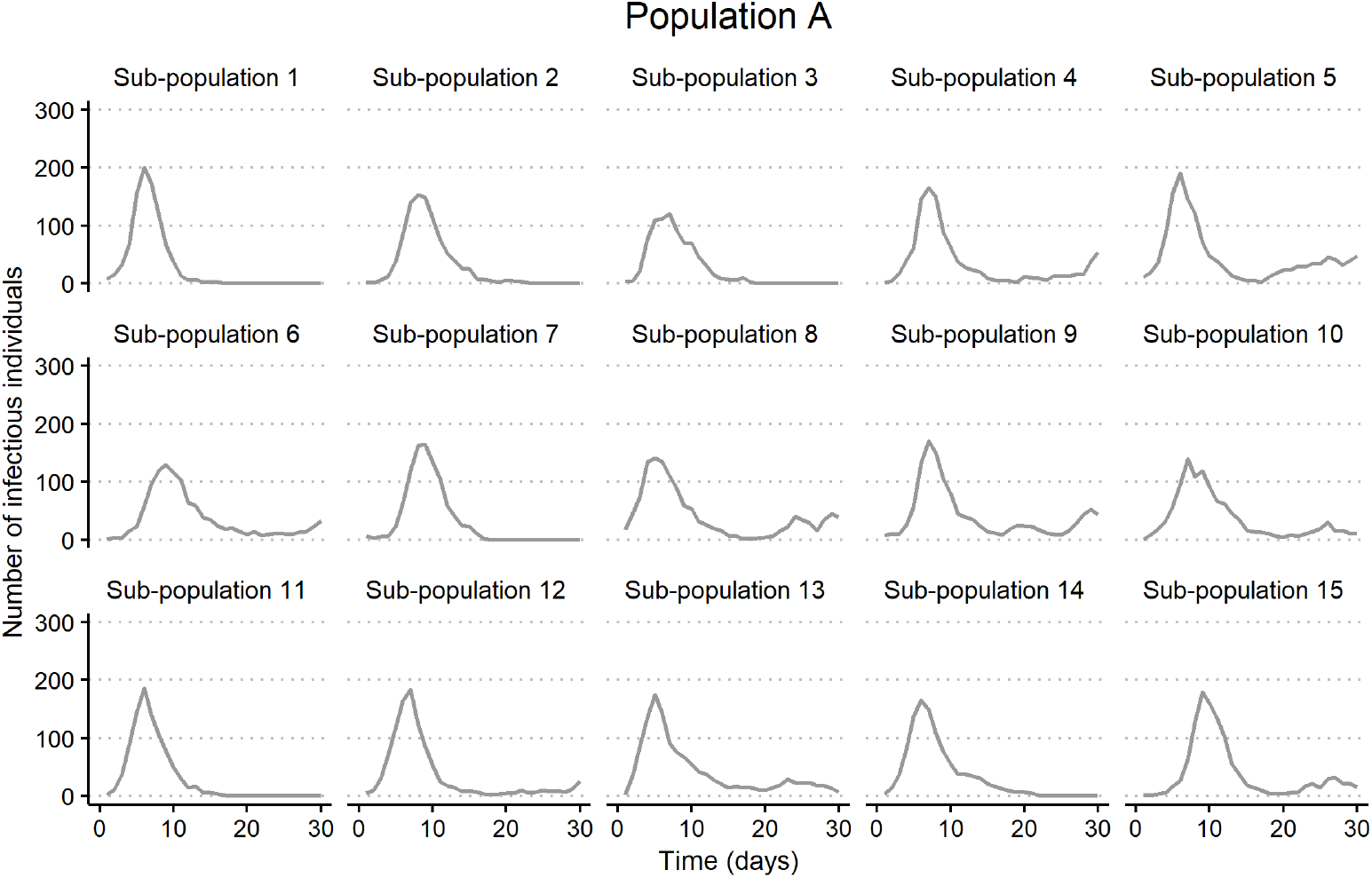
Synthetic dataset of Population A. Each panel shows the time-series data for a sub-population.

**Figure 3:**
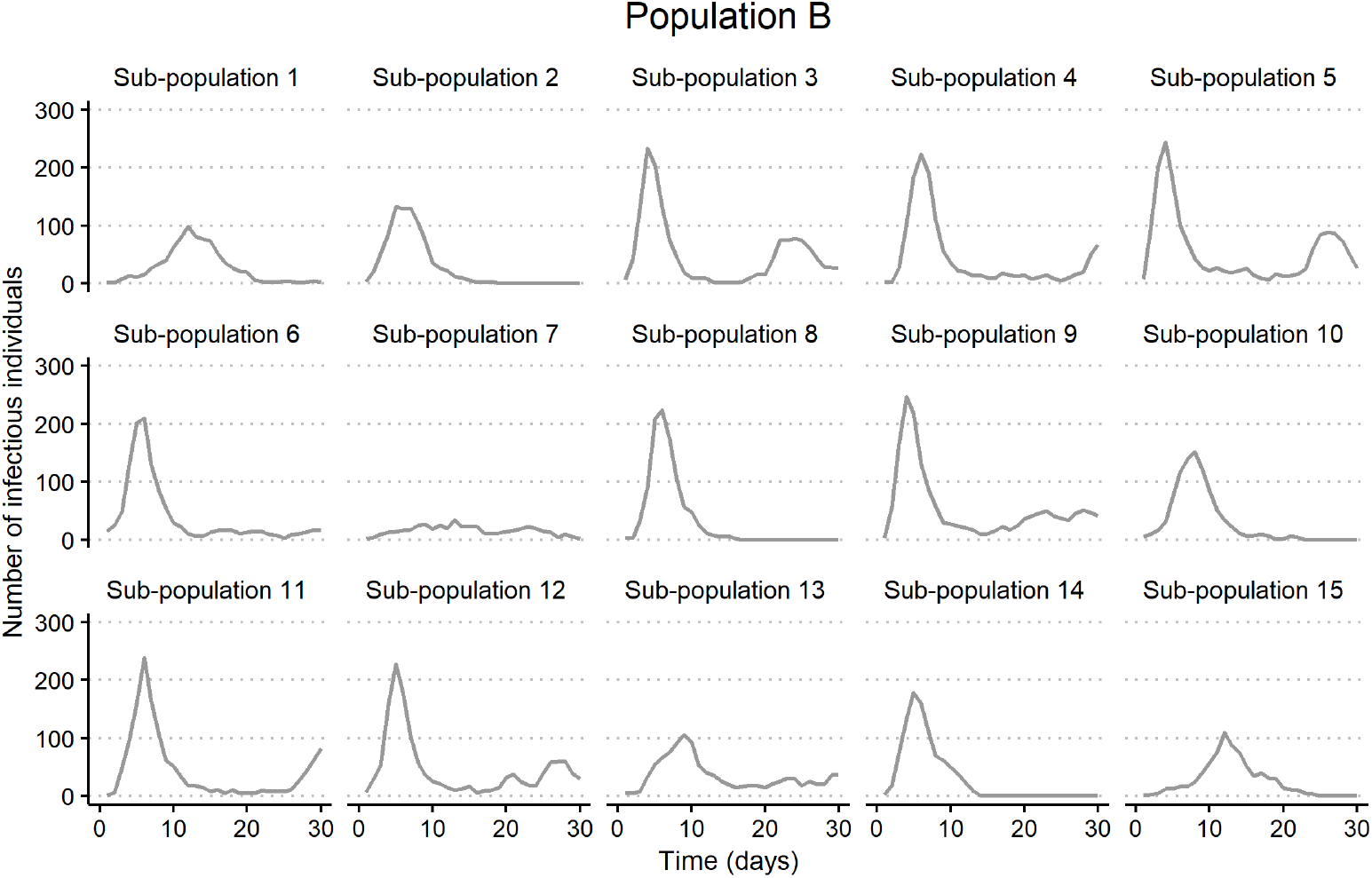
Synthetic dataset of Population B. Each panel shows the time-series data for a sub-population.

### 3.5 Further assumptions and modelling framework for parameter estimation for the two populations

To identify the presence and the magnitude of variability between the model parameters of the sub-populations in a population, the parameters of each of the sub-populations must be estimated. We started by assuming that both Populations A and B consisted of variable model parameters in the sub-populations and applied the two-step Bayesian hierarchical methodology described above. Our aim was to decide if there is enough evidence to support that variability in model parameters is present in the populations based on the two datasets.

We modelled each sub-population using the *SIRS* model described in Section 2.1. We fixed the recovery rate, *γ*, and the waning immunity rate, *µ* at their true values (1 and 0.06 respectively) across all the sub-populations in both populations and took the transmission rates *β*_*k*_ across the sub-populations as unknowns. Within a Bayesian hierarchical modelling structure, for the sub-population *k*, the observed infectious individuals consisted of the vector ***y***_***k***_ = (*I*_*k*_(1), *I*_*k*_(2), …, *I*_*k*_(30)) for all *k* = 1, …, 15 given *SIRS* model parameters (*β*_*k*_, *γ, µ*). The conditional prior of *β*_*k*_ was taken as a 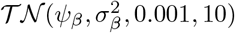. Here 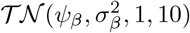 is a truncated normal distribution with mean *ψ*_*β*_, variance 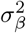, truncated to the (0.001, 10) interval. The hyper-prior distribution of *ψ*_*β*_ was uniform(0.001, 10) and the hyper-prior distribution of *σ*_*β*_ was uniform(0, 2.5).

To apply the two-step method to estimate the parameters, first, we used the ABC-SMC algorithm by Toni et al. (2009) in all the sub-populations independently. We took the prior distribution for all the sub-populations as a uniform distribution on (0.001, 10).

Within the ABC-SMC algorithm framework, we needed to choose a set of pre-defined tolerance values through a pre-defined number of generations. Across seven generations of the algorithm, the changing tolerance values that we used were (350, 300, 270, 250, 200, 170, 150). For sub-population 7 of dataset 2, we used (350, 300, 270, 250, 200, 150, 90) to avoid the particle system getting stuck on a local mode. Further details of this can be found in Supplementary Material S2. Furthermore, to implement the ABC-SMC algorithm by Toni et al. (2009), a kernel density, often known as a perturbation kernel, is also needed. A detailed explanation on choosing an appropriate perturbation kernel can be found in texts such as Filippi, Barnes, Cornebise, and Stumpf (2013). We used a normal distribution with an estimated variance calculated from the particles of the previous generation. For each sub-population, we obtained 5000 samples from the marginal posterior of *β*_*k*_. Next, these samples were fed into the MCMC algorithm (see Supplementary Material S1) to obtain 10000 samples from the marginal posteriors of *ψ*_*β*_ and *σ*_*β*_. After applying the MCMC step, as initial burn-in, we discarded the first 5000 iterations and we checked the convergence of the MCMC chains before obtaining the final samples of size 5000 each. Additional details, graphs, and diagnostics related to this MCMC step can be found in the Supplementary Material S2.

The second step was implemented using the ABC algorithm in Supplementary Material S1 with a tolerance value 150. For sub-population 7 of dataset 2, it was 90. We obtained 5000 samples from the posteriors of sub-population specific *β*_*k*_s for each sub-population under a hierarchical model. A visual comparison between the marginal posteriors obtained under a hierarchical and non-hierarchical model (where each sub-population is treated independently) can be found in the Supplementary Material S2. We implemented all the calculations in MATLAB (versions 2020a and 2020b) primarily using the Parallel Computing Toolbox across 32 virtual machines in the Nectar Research Cloud. All the codes that were used are publicly available. See Supplementary Material S5 for details.

## 4 Results

### 4.1 Parameter estimation

Figure 4 illustrates the marginal posterior distributions of *ψ*_*β*_ and *σβ*. Table 3 shows the point estimates for the posterior medians, the Highest Posterior Density (HPD) intervals and the width of the HPD intervals of the two populations. The HPD intervals were calculated using the Chen-Shao algorithm (Chen, Shao, & Ibrahim, 2012).These results validate that the first step of our two-step methodology to estimate the hyper-parameters is effective in the sense that the posterior medians are in close proximity with the true parameter values.

**Table 3:** Highest Posterior Density (HPD) intervals of the hyper-parameters

|  |  | Posterior median | Posterior mode | 95% HPDI | Width of the HPDI |
| --- | --- | --- | --- | --- | --- |
| $\psi_\beta$ | Population A | 1.9701 | 1.9494 | (1.8655, 2.0868) | 0.2213 |
|  | Population B | 2.1047 | 2.0511 | (1.8486, 2.4053) | 0.5567 |
| $\sigma_\beta$ | Population A | 0.0639 | 0.800 | (0.0002, 0.1823) | 0.1821 |
|  | Population B | 0.4971 | 0.2912 | (0.2768, 0.7694) | 0.4926 |

**Figure 4:**
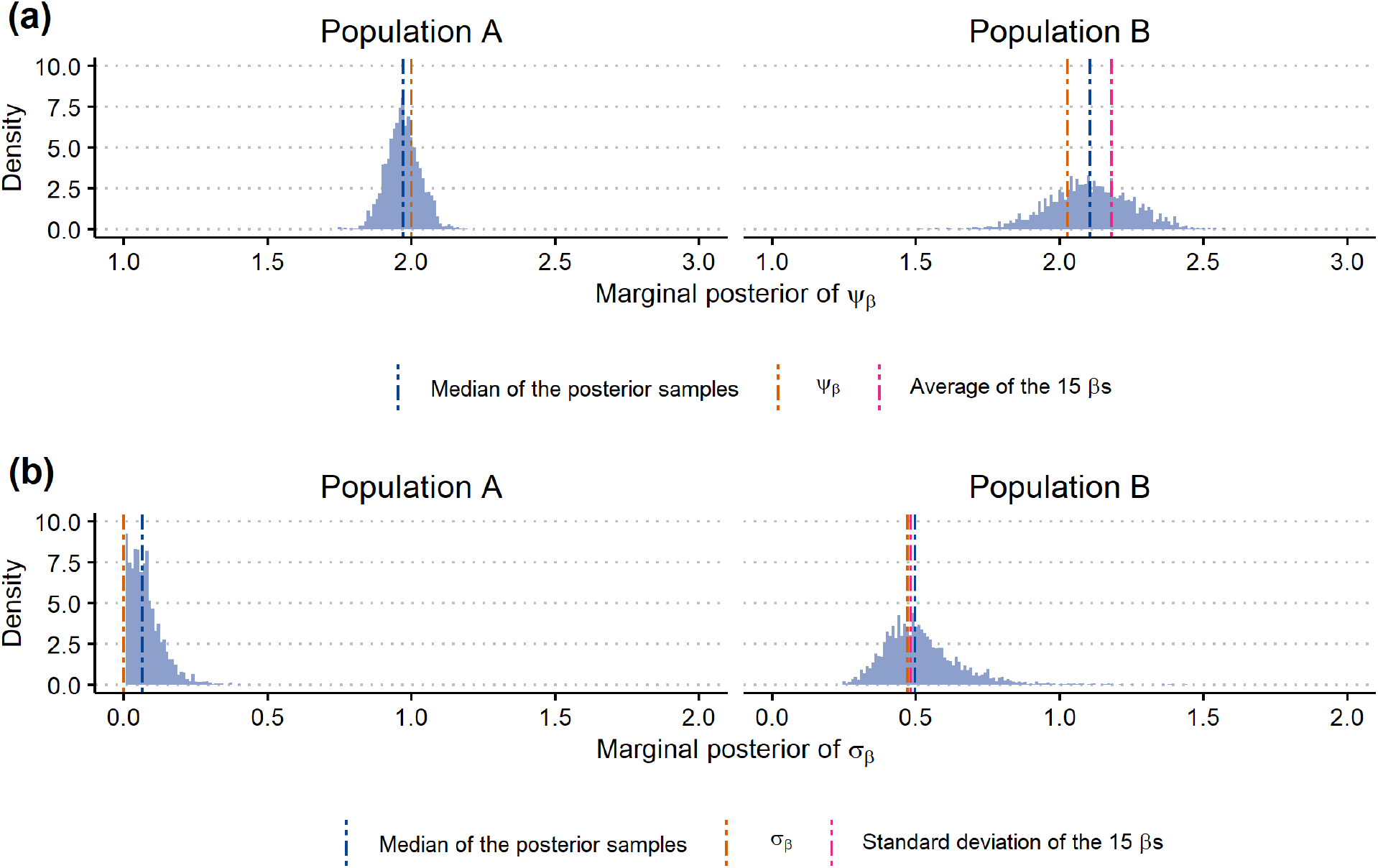
**(a)**: Marginal posterior of *ψ*_*β*_ for Populations A (left panel) and B (right panel). (b) : Marginal posterior of *σ*_*β*_ for Populations A (left panel) and B (right panel).

While both point estimates of *ψ*_*β*_ of the two populations are close to their true parameter values, the variance of the posterior distribution of Population A is smaller than that of Population B. The resulting HPD interval of population A is narrower. The marginal posteriors of standard deviation *σ*_*β*_ for both populations demonstrates that there is strong evidence that the variance of Population A is very small (almost negligible) in comparison to Population B, whose posterior median is equal to 0.4971. While the HPD interval for population A is (0.0002, 0.1823), for Population B, it is (0.2768, 0.7694).

Figure 5 illustrates the sub-population specific posteriors for *β*_*k*_s for the two populations under the hierarchical modelling framework. The posteriors of all the sub-populations in Populations A are visually similar, and the medians and 95% HPD intervals (see Supplementary Material S2) of the posterior are similar. The posteriors for the sub-population specific *β*_*k*_s of Population B are different in shape, and the posterior medians and 95% HPD intervals (see Supplementary Material S2) show marked variation.

**Figure 5:**
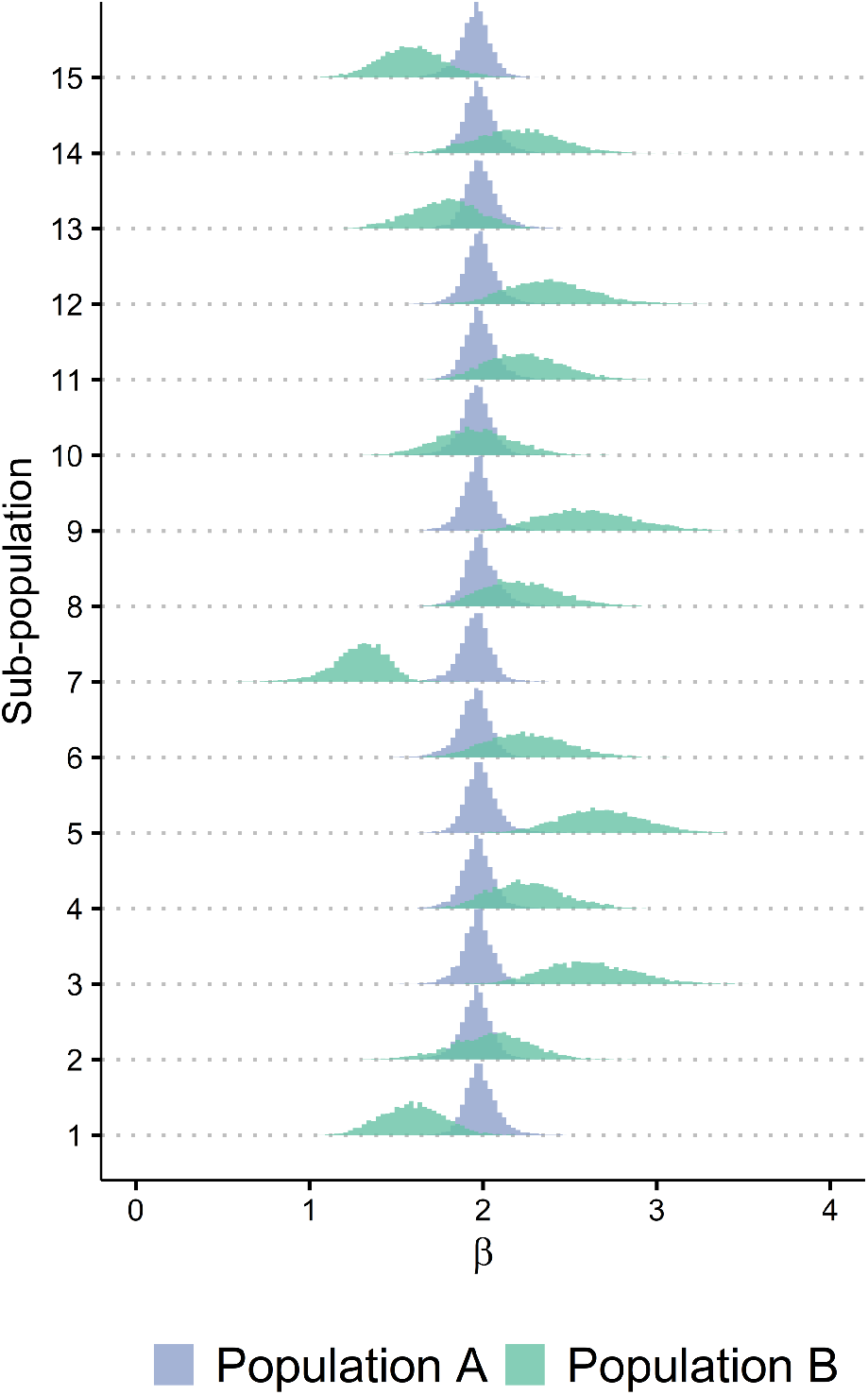
Marginal posteriors of *β*_*k*_ (for *k* = 1, 2, …, 15) for Populations A and B.

We have demonstrated that under a hierarchical modelling framework, we can accurately estimate parameters using ABC methods using our two-step algorithmic set-up. The presence of variability between model parameters can be identified by visual diagnostics of the marginal posterior distributions of the *ψ*_*β*_, *σ*_*β*_ (hyper-parametric level) as well from the sub-population specific posteriors of *β*_*k*_. Further evidence of the presence of variability between model parameters can be clarified using the Region of Practical Equivalence (ROPE) (Kruschke, 2013) criterion (refer to Supplementary Material S2).

### 4.2 Estimating the probability of epidemic fade-out

In order to identify the affects of estimated model parameters on the probability of epidemic fade-out at both the population and sub-population level, we calculated the probability of epidemic fade-out with estimated transmission rates, recovery rate of 1, and waning immunity rate of 0.06 using the algorithms implemented by Ballard et al. (2016). Refer to the Supplementary Material S3 for further details about this implementation.

We considered an *SIRS* model with unknown transmission rate *ψ*_*β*_, recovery rate *γ* = 1 and waning immunity rate *µ* = 0.06 for a sub-population of size 1000, and with initially one infectious individual. Then the epidemic fade-out probabilities were calculated under each sampled value of the posterior of *ψ*_*β*_. Refer to Figure 6 and Table 4 for the change of fade-out probability with respect to *ψ*_*β*_ and the distributions of the fade-out probabilities for Populations A and B. For Population A, the median probability of epidemic fade-out is 0.5197 and that of Population B is 0.48098. The variability of epidemic fade-out probabilities in Population A is smaller than that of Population B.

**Table 4:** Epidemic fade-out probabilities under *ψ*_*β*_

| Table 4: Epidemic fade-out probabilities under $\psi_\beta$ | | | |
| --- | --- | --- | --- |
| Population A | 95% HPD lower level | Posterior median | 95% HPD upper level |
| $\psi_\beta$ | 1.8655 | 1.9701 | 2.0868 |
| Fade-out probability | 0.5505 | 0.51797 | 0.4866 |
| Population B | 95% HPD lower level | Posterior median | 95% HPD upper level |
| $\psi_\beta$ | 1.8486 | 2.1047 | 2.4053 |
| Fade-out probability | 0.5521 | 0.48098 | 0.4053 |

**Figure 6:**
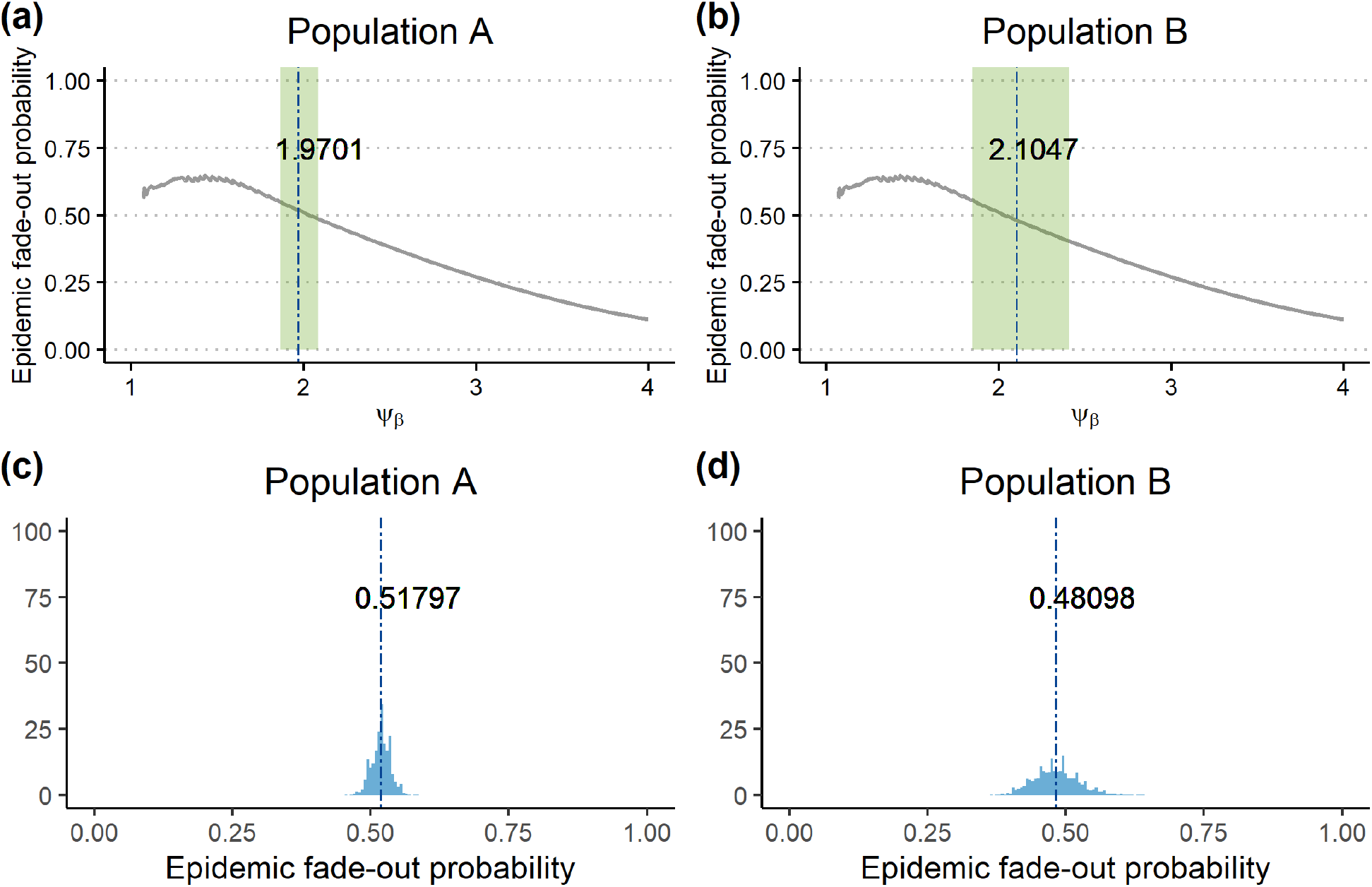
Change of probability of epidemic fade-out with respect to *ψ*_*β*_ in grey are shown in panels (a) and (b). Green solid areas are the 95% HPD intervals of the posterior of *ψ*_*β*_ under each population. The dark blue dashed lines are the posterior medians. The distribution of the estimated fade-out probabilities using the estimated marginal posteriors of *ψ*_*β*_ of the two populations are given in light blue in (c) and (d) panels. The dark blue dashed lines are median probabilities of epidemic fade-out.

The distributions of the estimated epidemic fade-out probabilities of the sub-populations of Populations A and B are illustrated in Figure 7. While the distributions of Population A looked similar and had a median value close to 0.52, the distributions of Population B were different and their medians had significant variability. In Population B, the distributions of the fade-out probabilities of sub-populations 1, 7, 13, and 15 are skewed distributed with a median fade-out probability around 0.6. The posterior medians of *β*_*k*_ of these sub-populations were close to 1.5 where the probability of epidemic fade-out is at its peak when the recovery rate and waning immunity rate are 1 and 0.06 respectively. Furthermore, given the estimated posterior samples from the marginals of *β*_*k*_, the ranges that covered the epidemic fade-out probabilities of all these sub-populations were distinctly different (see Supplementary Material S3). While sub-populations 1, 7, 13 and 15 had a considerably high probability of epidemic fade-out, sub-populations 3, 5 and 9 had a rather small median epidemic fade-out probabilities (see Supplementary Material S3).

**Figure 7:**
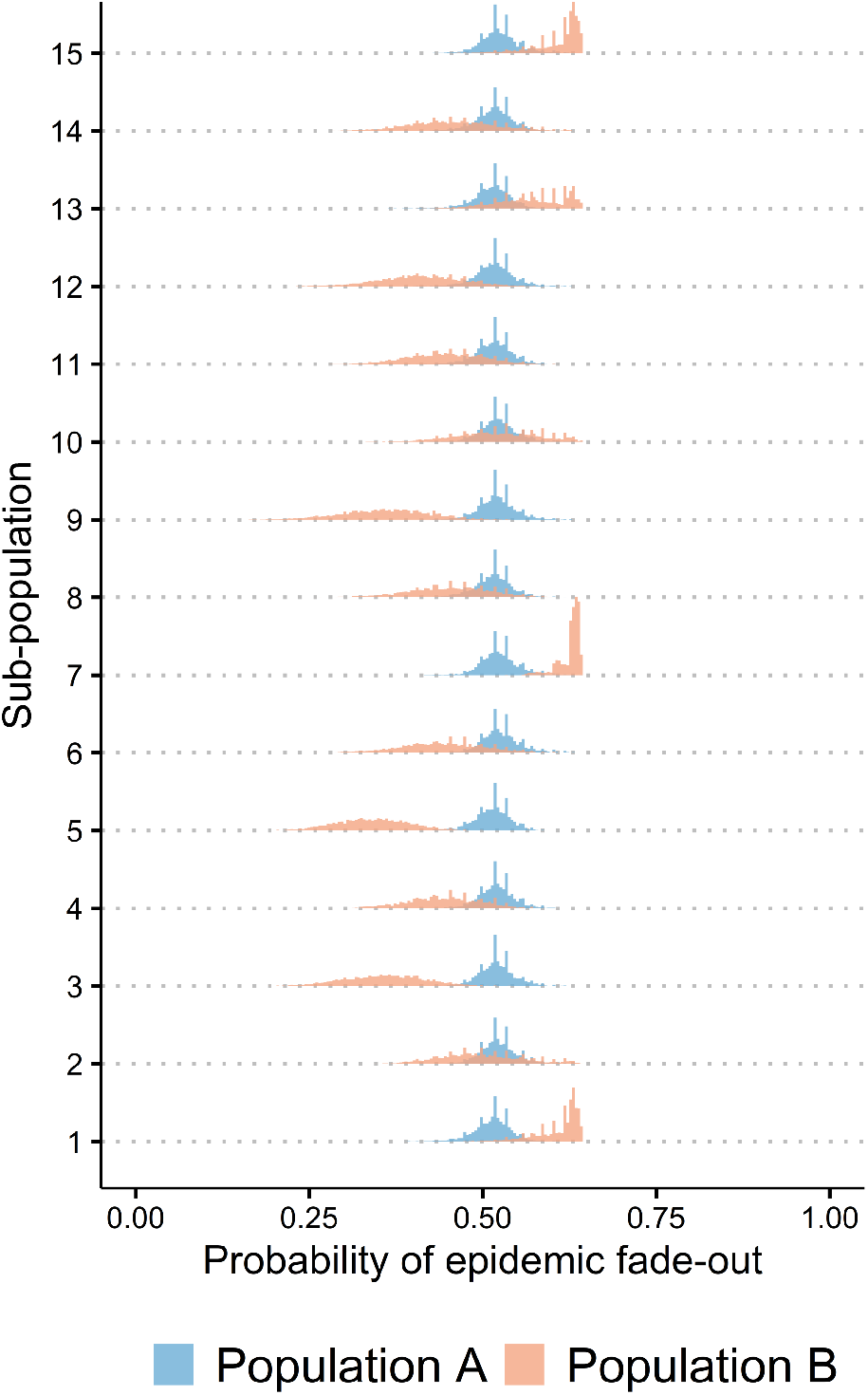
The distributions of the probability of epidemic fade-out of the sub-populations of Populations A and B.

### 4.3 Suitability of using a hierarchical modelling framework for multiple outbreak data

We observed that while the time series data of sub-population 7 was visually different from the other sub-populations, its true parameter value, *β*_7_, was also significantly different and yet, our estimation framework yielded accurate results. Motivated by this observation, we were interested in examining the impact of using a hierarchical modelling framework under extreme data conditions. Therefore, we implemented the following simulation experiment.

We generated synthetic data for another (i.e, 16th) sub-population for Population B using the same synthetic data generation process we described in subsection 3.4. We sampled an extreme value for transmission rate, *β*_16_ = 3.2516 and generated a sample path with a higher peak than the others (i.e., visually different than others). The true extinction probability for this sub-population is 0.2218. We then carried out parameter estimation for all the sub-populations independently without implementing a hierarchical structure (using the ABC-SMC algorithm by Toni et al. (2009)) and adding a hierarchical modelling framework using our two-step methodology with the additional modelling conditions described in the subsection 3.5. We further estimated the probabilities of epidemic fade-out using the same methodology we described in subsection 4.2 using the *β*_16_ that were estimated both independently and under a hierarchical modelling framework. See Figure 8 for the results obtained from this experiment.

**Figure 8:**
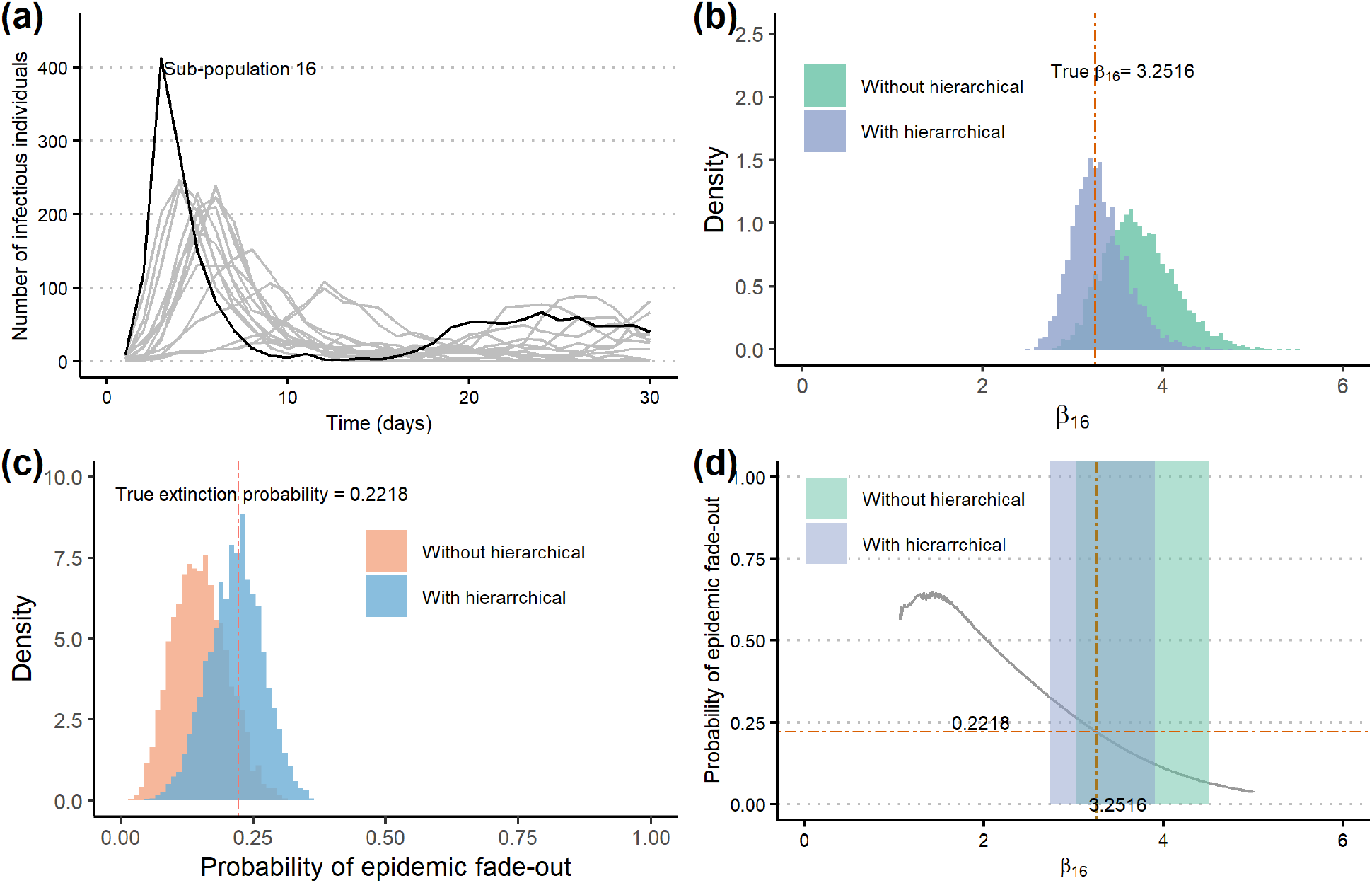
**(a):** Time series data of sub-population 16 (in black). Time series data of other sub-populations in Population B are presented in grey. **(b) :** Comparison of the marginal posterior of *β*_16_ with (in purple) and without (in green) a hierarchical modelling framework. Posterior median under a hierarchical modelling framework is closer to *β*_16_. **(c):** Comparison of the epidemic fade-out probabilities estimated with (in blue) and without (in pink) a hierarchical model.Median extinction probability estimated under a hierarchical model is closer to the true extinction probability. **(d) :** Change of probability of epidemic fade-out with *β* (in grey). The 95% HPD interval for *β*_16_, (3.0260, 4.5087), without the hierarchical model is shown in green and that of the hierarchical model, (2.7291, 3.8911), is shown in purple. The brown dashed line is the true *β*_16_ value, 3.2516. The orange dashed line is the true epidemic fade-out probability, 0.2218, of the 16th sub-population

We noticed that even when data and the true parameters can be as extreme as in sub-population 16, using a hierarchical modelling framework can yield more accurate estimates. Our results showed that estimates for the parameter, *β*_16_, and the epidemic fade-out probability improved when estimation was done with a hierarchical modelling framework.

## 5 Discussion

We have shown that parameter estimation carried out with a Bayesian perspective using our new two-step methodology, coupled with ABC methods, is a suitable and efficient approach to estimate parameters of hierarchical epidemic models.

We have shown that hierarchical modelling is a powerful approach and we can accurately estimate not only the sub-population specific parameters but also learn about the common distribution that the model parameters of the sub-populations are sampled from.

By estimating the parameters of the two synthetic datasets using Bayesian hierarchical methods, we have demonstrated that we can identify the presence of variability between model parameters of the sub-populations. If we infer that a population consisted of sub-populations with identical model parameters, we can reasonably attribute the different observed sample paths to be a result of stochastic variations. On the other hand, if we infer that a population consisted of variable model parameters, different sample paths of the sub-populations can be better explained by models with sub-population specific parameters as well as stochastic effects.

As a result of using hierarchical parameter estimation techniques, we can study epidemic fade-out probabilities under two levels. One is to make inferences on each sub-population’s probability of epidemic fade-out and conclude if epidemic fade-outs were observed, whether it was due to stochastic fluctuations only or else, events attributed by the estimated model parameters. The next level is making use of the inference made at the population level. Based on the inference made on the hyper-parameters, it is possible to predict the probability of epidemic fade-out when an epidemic breaks out in another sub-population of the general population.

Finally, when outbreak data are present in multiple sub-populations, we have illustrated that the use of a hierarchical modelling approach to estimate parameters can produce more accurate results in comparison to estimating the parameters independently. We further showed that estimated epidemic fade-out probabilities are also accurate when the estimation was approached via hierarchical modelling methods.

We have considered that data were perfectly observed, However, this is not realistic. Generally, data are often distorted by factors such as under/ over-reporting, misdiagnosis, equipment error. Epidemic fade-outs may not be completely observed as well. Therefore, it would be ideal to apply our methods to other simulation experiments or real data under different settings. External intervention measures may also be present while data are being observed and thereupon, it is noteworthy to study data under such circumstances. We also have considered that the transmission rate of a sub-population is the only unknown parameter. This is not true in practice. For example, by considering both the recovery rate and the waning immunity rate as unknown factors, we would have to consider a higher dimensional statistical analysis that would be suitable for not only an *SIRS* model with three unknown model parameters but also for similar complex epidemic models.

Thus far, we have applied this framework using *SIRS* models, with an emphasis on the epidemic fade-out that can be observed in multiple sub-populations. This framework is applicable and generalisable to other biological processes in which average dynamics display damped oscillatory behaviour with a deep initial trough. For instance, some malaria-infected individuals go through recrudescence of the infection while others do not (examples: Cao et al. (2019); Collins et al. (2018)), in ecology where some species become extinct and others do not, and in bacterial populations where heterogeneous conditions may or may not cause extinction.

## 6 Acknowledgements

The authors would like to thank Peter Ballard for his support and helpful discussions in understanding his codes in Python to estimate the probabilities of epidemic fade-out as in his paper Ballard et al. (2016). All the MATLAB computations were carried out by the use of the Nectar Research Cloud (project *Infectious Diseases*), a collaborative Australian research platform supported by the National Collaborative Research Infrastructure Strategy (NCRIS).

## Notes

### Competing Interest Statement

The authors have declared no competing interest.

