## Supplementary Material for "Estimation of the probability of epidemic fade-out from multiple outbreak data"

#### S1 Hierarchical parameter estimation: theory and algorithms

##### S1.1 Step 1: Estimation of hyper-parameters

In this step, we will assume that we have done part (a) of Step 1 and that we have $N_1$ samples from the marginal posteriors of each sub-population. During this step, we propose to use a common non-informative prior distribution $p(\boldsymbol{\theta}_k)$ across all the $K$ sub-populations that cover the parameter space of the prior distribution (hyper-prior) of the mean parameter of the hyper-distribution. This will ensure that the parameter space of the conditional prior $p(\boldsymbol{\theta}_k|\psi)$ will be explored within the hierarchical framework.

We will focus on implementing part (b) of Step 1. In this particular part, in order to implement the MCMC algorithm at the hyper-parameter level, we first need to define an estimate to the likelihood for all the sub-populations. This section is reserved for this purpose.

The marginal posterior distribution for hyper-parameters can be obtained by integrating out the population specific parameters such that,

$$p(\boldsymbol{\psi}|\boldsymbol{y}) = \frac{\left[\prod_{k=1}^{K} \int_{\boldsymbol{\theta}_k} p(\boldsymbol{y}_k|\boldsymbol{\theta}_k)p(\boldsymbol{\theta}_k|\boldsymbol{\psi})d\boldsymbol{\theta}_k\right] p(\boldsymbol{\psi})}{p(\boldsymbol{y})} \tag{1}$$

$$= \frac{[\prod_{k=1}^{K} p(\boldsymbol{y_k}|\boldsymbol{\psi})]p(\boldsymbol{\psi})}{p(\boldsymbol{y})},$$

where,

$$p(\boldsymbol{y_k}|\boldsymbol{\psi}) = \int_{\boldsymbol{\theta_k}} p(\boldsymbol{y}_k|\boldsymbol{\theta}_k)p(\boldsymbol{\theta}_k|\boldsymbol{\psi})d\boldsymbol{\theta}_k. \tag{2}$$

Noting that $p(\boldsymbol{y})$ is a constant,

$$p(\boldsymbol{\psi}|\boldsymbol{y}) \propto \left[\prod_{k=1}^{K} p(\boldsymbol{y}_k|\boldsymbol{\psi})\right] p(\boldsymbol{\psi}). \tag{3}$$

First we will consider one sub-population, say the $k$th sub-population. Let the posterior distribution for this sub-population be denoted by $p(\boldsymbol{\theta_k}|\boldsymbol{y}_k)$ and assume that it can be approximated by the distribution from which we sample parameter values from using parameter estimation algorithm such as an ABC algorithm implemented for the $k$th sub-population only. Let this approximated posterior be denoted as $q(\boldsymbol{\theta_k}|\boldsymbol{y}_k)$ and therefore,

$$q(\boldsymbol{\theta_k}|\boldsymbol{y}_k) \approx p(\boldsymbol{\theta_k}|\boldsymbol{y}_k),$$

*

and

$$\boldsymbol{\theta_k}^{(j)} \sim q(\boldsymbol{\theta_k}|\boldsymbol{y}_k),$$

where $j = 1, 2, ..., N_1$ and $N_1$ is the size of the accepted sample from the marginal posteriors of the $k$th sub-population specific parameters. Let the pool of accepted sample from the marginal posteriors of the $k$th sub-population specific parameters from an ABC based algorithm be $\{\boldsymbol{\theta}_k^{(j)}\}_{j=1}^{N_1}$.

Now, we want to be able to compute 2 using $\{\boldsymbol{\theta}_k^{(j)}\}_{j=1}^{N_1}$. In order to computationally enable this, next, we will consider using importance sampling theory to approximate the integration. (Use of importance sampling theory to approximate this integration has also been proposed by Wu, Angelikopoulos, Beck, and Koumoutsakos (2019) to calculate estimates for hyper-parameters with models that have tractable likelihoods with similar methods to this). This approximated integration can then be used as an estimate in an MCMC acceptance rejection criteria.

Suppose we use $q(\boldsymbol{\theta_k}|\boldsymbol{y}_k)$ as the importance sampling distribution and $\{\boldsymbol{\theta_k}^{(j)}\}_{j=1}^{N_1}$ as the sampled values from it. Therefore, using 2,

$$\begin{aligned} p(\boldsymbol{y_k}|\boldsymbol{\psi}) &= \int_{\boldsymbol{\theta_k}} p(\boldsymbol{y}_k|\boldsymbol{\theta}_k)p(\boldsymbol{\theta}_k|\boldsymbol{\psi})d\boldsymbol{\theta}_k \\ &= \int_{\boldsymbol{\theta_k}} \frac{p(\boldsymbol{y}_k|\boldsymbol{\theta}_k)p(\boldsymbol{\theta}_k|\boldsymbol{\psi})}{q(\boldsymbol{\theta_k}|\boldsymbol{y}_k)} q(\boldsymbol{\theta_k}|\boldsymbol{y}_k)d\boldsymbol{\theta}_k. \end{aligned} \tag{4}$$

Expression 4 can be approximated by using $\{\boldsymbol{\theta}_k^{(j)}\}_{j=1}^{N_1}$ as,

$$\begin{aligned} p(\boldsymbol{y_k}|\boldsymbol{\psi}) &\approx \frac{1}{N_1}\sum_{j=1}^{N_1} \frac{p(\boldsymbol{y}_k|\boldsymbol{\theta}_k^{(j)})p(\boldsymbol{\theta}_k^{(j)}|\boldsymbol{\psi})}{q(\boldsymbol{\theta_k^{(j)}}|\boldsymbol{y}_k)} \\ &\approx \frac{1}{N_1}\sum_{j=1}^{N_1} \frac{p(\boldsymbol{y}_k|\boldsymbol{\theta}_k^{(j)})p(\boldsymbol{\theta}_k^{(j)}|\boldsymbol{\psi})p(\boldsymbol{y}_k)}{p(\boldsymbol{y}_k|\boldsymbol{\theta_k^{(j)}})p(\boldsymbol{\theta_k^{(j)}})} \\ &\approx \frac{p(\boldsymbol{y}_k)}{N_1}\left[\sum_{j=1}^{N_1} \frac{p(\boldsymbol{\theta}_k^{(j)}|\boldsymbol{\psi})}{p(\boldsymbol{\theta}_k^{(j)})}\right], \end{aligned} \tag{5}$$

where $p(.|\boldsymbol{\psi})$ is the hyper-distribution whose functional form is considered to be known (usually a normal distribution with unknown hyper-parameters) and $p(\boldsymbol{\theta}_k)$ is the prior distribution of the $k$th population used in the ABC algorithm (in Step 1 (a)) to obtain the sample of $\{\boldsymbol{\theta_k}^{(j)}\}_{j=1}^{N_1}$.

Now, we consider an approximate value for the overall likelihood of data of all the populations such that,

$$\begin{aligned} p(\boldsymbol{y}|\boldsymbol{\psi}) &\approx \prod_{k=1}^{K} \frac{p(\boldsymbol{y}_k)}{N_1}\left[\sum_{j=1}^{N_1} \frac{p(\boldsymbol{\theta}_k^{(j)}|\boldsymbol{\psi})}{p(\boldsymbol{\theta}_k^{(j)})}\right] \\ &= \frac{1}{N_1}\prod_{k=1}^{K}\sum_{j=1}^{N_1} \frac{p(\boldsymbol{\theta}_k^{(j)}|\boldsymbol{\psi})p(\boldsymbol{y}_k)}{p(\boldsymbol{\theta}_k^{(j)})} = \hat{p}(\boldsymbol{y}|\boldsymbol{\psi}). \end{aligned} \tag{6}$$

Noting that $N_1$ and $p(\boldsymbol{y_k})$ are constants with respect to each sub-population,

$$\hat{p}(\boldsymbol{y}|\boldsymbol{\psi}) \propto \prod_{k=1}^{K}\sum_{j=1}^{N_1} \frac{p(\boldsymbol{\theta}_k^{(j)}|\boldsymbol{\psi})}{p(\boldsymbol{\theta}_k^{(j)})}. \tag{7}$$

The expression 3 is now proportional to

$$p(\boldsymbol{\psi}|\boldsymbol{y}) \propto \hat{p}(\boldsymbol{y}|\boldsymbol{\psi})p(\boldsymbol{\psi}). \tag{8}$$

Now that we have a simplified form for the marginal posterior distribution of the hyper-parameters, we can now easily use 8 in the acceptance rejection criterion for the $b$ th iteration of an MCMC, Metropolis-Hastings algorithm such that,

$$\alpha = min(1, \frac{\hat{p}(\boldsymbol{y}|\boldsymbol{\psi^{can}})p(\boldsymbol{\psi^{can}})q(\boldsymbol{\psi}^{(b-1)}|\boldsymbol{\psi}^{can})}{\hat{p}(\boldsymbol{y}|\boldsymbol{\psi^{(b-1)}})p(\boldsymbol{\psi}^{(b-1)})q(\boldsymbol{\psi}^{can}|\boldsymbol{\psi}^{(b-1)})}), \tag{9}$$

where $\boldsymbol{\psi}^{can}$ is the proposed hyper-parameter value from the $q(.)$ proposal distribution. Using a functional form for the proposal $q(.)$ similar to that of $p(.|\boldsymbol{\psi})$ ensures efficiency in the sense that the explored parameter space from the random walk is controlled. The full description of the algorithm is presented in 1.

If the prior distributions $p(\boldsymbol{\theta}_k^{can})$ (when the hierarchical structure is disregarded) are selected to be uniform distributions with the same intervals for all the populations, then the values of $p(\boldsymbol{\theta}_k^{can})$ are equally likely and therefore, the importance sampling estimate can be further simplified to,

$$\hat{p}(\boldsymbol{y}|\boldsymbol{\psi}) \propto \prod_{k=1}^{K}\sum_{j=1}^{N} p(\boldsymbol{\theta}_k^{(j)}|\boldsymbol{\psi}). \tag{10}$$

It must be examined if $\boldsymbol{\theta_k^{(j)}}$ is a possible sample from $p(.|\boldsymbol{\psi})$ for all the populations (that is $p(\boldsymbol{\theta}_k^{(j)}|\boldsymbol{\psi})$ is non-zero). Algorithm 1 shows the final Metropolis-Hastings type procedure to estimate hyper-parameters.

**Algorithm 1 Step 1:** Hyper-parameter estimation for hierarchical data

1: **Inputs:** $\{\theta_k^{(j)}\}_{j=1}^{N_1}$, and $\boldsymbol{y}_k$ for all $k = 1, ..., K$ and $j = 1, ...., N_1$
2: **Output:** $\{\boldsymbol{\psi}^{(b)}\}_{b=n}^{N_2}$, a sample from the marginal posteriors of the hyper-parameters after an initial burn-in period $b = 1, .., n$.
3: **Initialisation:** Define a hyper-prior distribution $p(\boldsymbol{\psi})$ and a proposal distribution $q(.)$. Set an initial value for the hyper-parameters as $\boldsymbol{\psi}^{(1)}$. Set the number of iterations to run as $N_2$.
4: **for** $b = 2 \ldots$ **do**
5: Propose a hyper-parameter $\boldsymbol{\psi}^{can}$ such that

$$\boldsymbol{\psi}^{can} \sim q(.|\boldsymbol{\psi}^{(b-1)})$$

6: Calculate the acceptance / rejection probability $\alpha$ as in 9.
7: Generate $u \sim uniform(0, 1)$
8: **if** $\alpha > u$ **then**
9: $\boldsymbol{\psi}^{(b)} = \boldsymbol{\psi}^{can}$
10: **else**
11: $\boldsymbol{\psi}^{(b)} = \boldsymbol{\psi}^{(b-1)}$
12: Continue Step 5 until convergence can be ensured.

**Note:**
See Supplementary Material S2.2 for some diagnostics that can be used to check convergence.

##### S1.2 Step 2: Estimation of sub-population specific parameters

Once samples from the marginal posteriors of the hyper-parameters are obtained, the second step of the algorithm can be carried out as in Algorithm 2. Since the sub-population specific parameters that are proposed in an ABC iteration are already conditioned and controlled by the hyper-parameters, convergence to the posteriors can easily be achieved without having to resort to more complex ABC-based algorithms, and using a basic ABC algorithm under an appropriate tolerance value (that can already be found through the first step) is sufficient.

**Algorithm 2 Step 2:** Sub-population specific parameter estimation under the hierarchical model

**Inputs:** $\{\boldsymbol{\psi}^{(b)}\}_{b=1}^{N_2}$
**Outputs:** $\{\boldsymbol{\theta}_k^{'(b)}\}_{b=1}^{N_2}$, for all $k = 1, ..., K$ and $b = 1, ...., N_2$; samples from sub-population specific posteriors under a hierarchical structure.
**for** k=1,2,. . . , K **do**
  **for** $b = 1, ..., N_2$ **do**
    **repeat**
      Generate $\boldsymbol{\theta}''_k$ from the conditional prior distribution $p(.|\boldsymbol{\psi}^{(b)})$
      Generate $\boldsymbol{x}$ from the likelihood $f(.|\boldsymbol{\theta}''_k)$
    **until** $\rho\{\eta(\boldsymbol{x}), \eta(\boldsymbol{y})\} \leq \epsilon$
    set $\boldsymbol{\theta}_k^{'(b)} = \boldsymbol{\theta}''_k$

### S2 Parameter estimation: Application

#### S2.1 Choosing tolerance values

Basic ABC algorithms were run under a very large tolerance value (400), to identify potential tolerance values that may lead the particle system being stuck on local modes. Figures S1 and S2 show 50000 samples/particles obtained for each sub-population with respect to the distance metric. Potential tolerance values (that are less than 400) that may lead to being caught in a local mode are displayed as vertical lines. Tolerance values for the ABC SMC algorithm was chosen by avoiding these particular values.

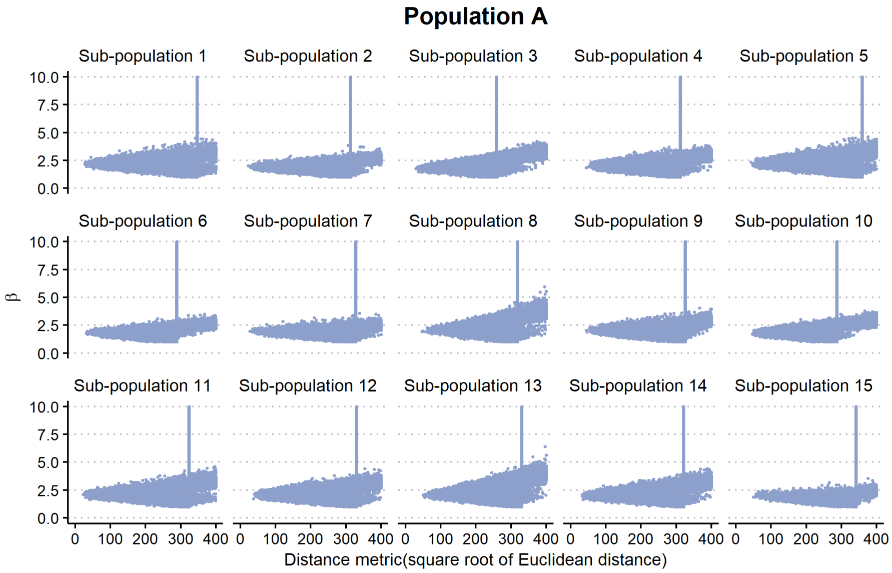


Figure S1: Scatter plots of $\beta$ vs distance metric for Population A

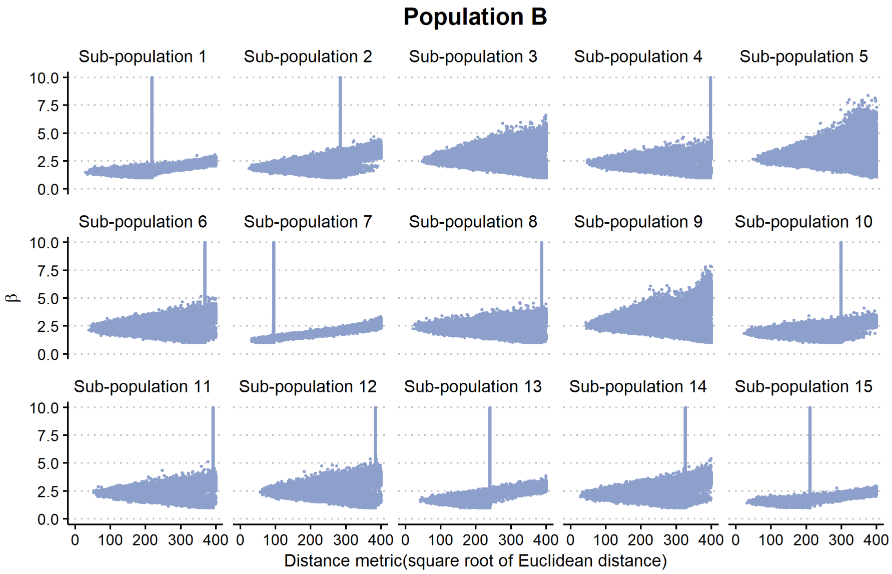


Figure S2: Scatter plots of $\beta$ vs distance metric for Population B

##### S2.2 Diagnostics: Step 1 of the two-step methodology to estimate hyper-parameters

**MCMC chains under different initial starting values:**

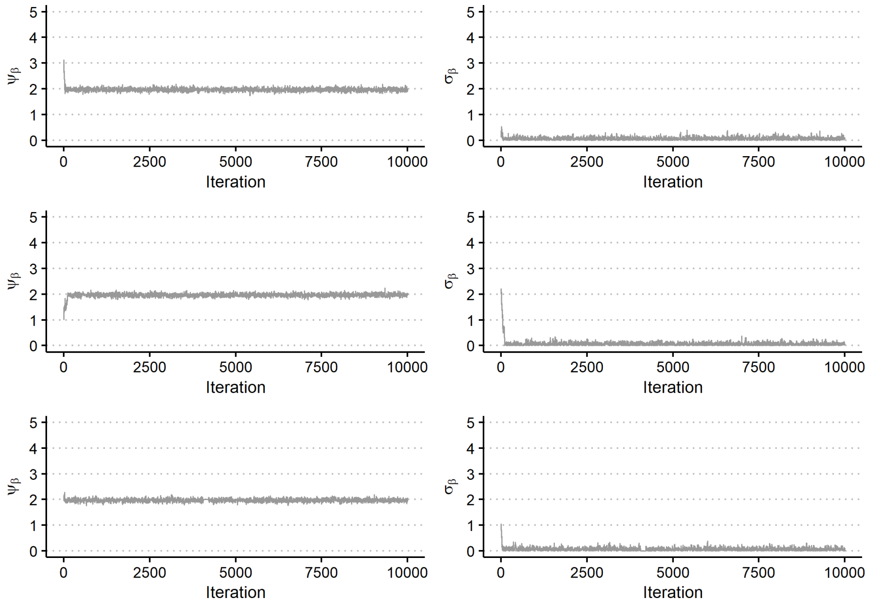


Figure S3: Population A: Each row is a different MCMC run with different initial $\psi_\beta$ and $\sigma_\beta$.

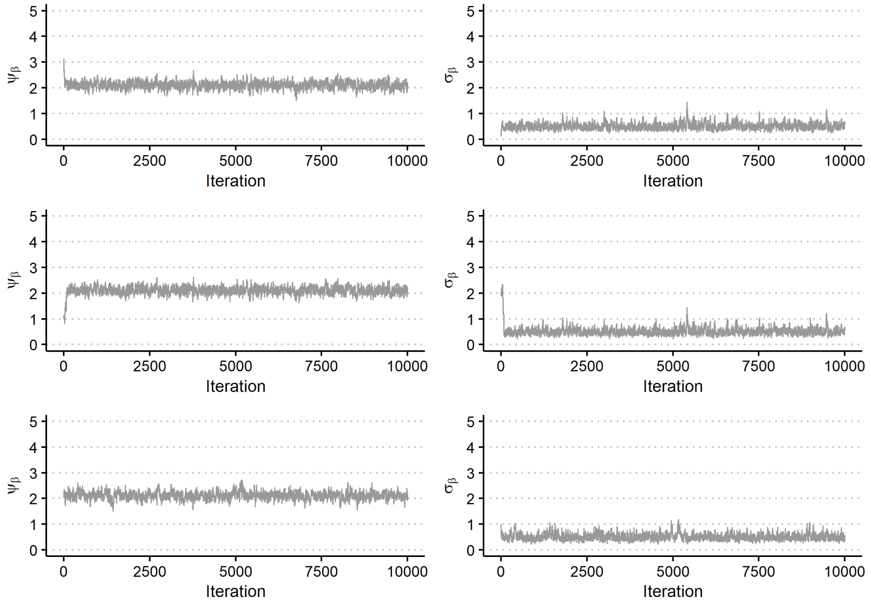


Figure S4: Population B: Each row is a different MCMC run with different initial $\psi_\beta$ and $\sigma_\beta$.

Proposal distributions for the $b$th iteration of an MCMC chain are

$$\psi_\beta^{(b)} \sim Normal(\psi_\beta^{(b-1)}, 0.1^2) \tag{11}$$

$$\sigma_\beta^{(b)} \sim Normal(\sigma_\beta^{(b-1)}, 0.1^2). \tag{12}$$

##### S2.3 Comparison of sub-populations specific parameters under a hierarchical model (Step 2 of the two-step methodology) and independently estimated values.

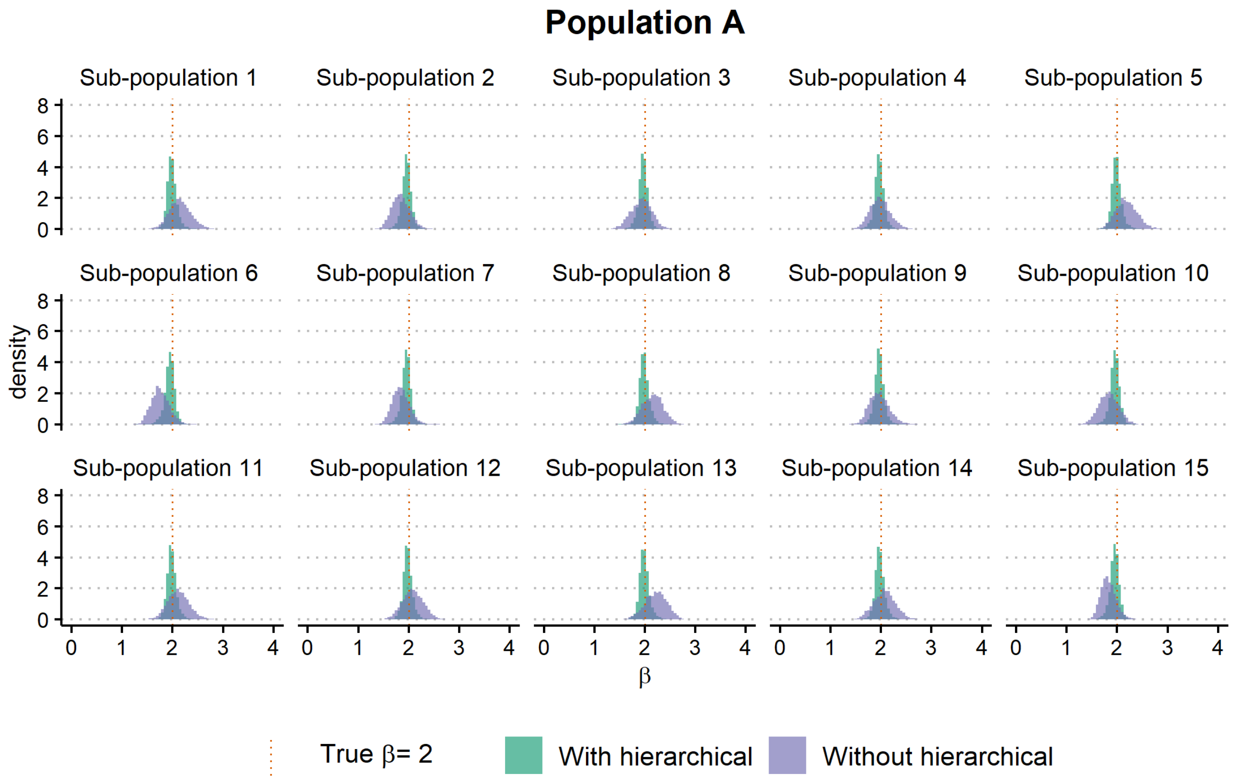


Figure S5: Marginal posteriors of $\beta$ for Population A

Table S1: Population A HPDS

| | Without a hierarchical model | | | | With a hierarchical model | | | |
|---|---|---|---|---|---|---|---|---|
| **Sub-population** | **HPD lower level** | **HPD upper level** | **Width of the HPD** | **Posterior median** | **HPD lower level** | **HPD upper level** | **Width of the HPD** | **Posterior median** |
| 1 | 1.7545 | 2.5860 | 0.8315 | 2.1414 | 1.8101 | 2.1861 | 0.3759 | 1.9806 |
| 2 | 1.4864 | 2.1742 | 0.6878 | 1.8380 | 1.7674 | 2.1497 | 0.3823 | 1.9607 |
| 3 | 1.5258 | 2.3013 | 0.7754 | 1.9360 | 1.7750 | 2.1522 | 0.3772 | 1.9681 |
| 4 | 1.5980 | 2.3696 | 0.7716 | 1.9761 | 1.7607 | 2.1561 | 0.3955 | 1.9662 |
| 5 | 1.7730 | 2.5863 | 0.8133 | 2.1786 | 1.7986 | 2.1920 | 0.3933 | 1.9826 |
| 6 | 1.4379 | 2.1085 | 0.6705 | 1.7454 | 1.7429 | 2.1387 | 0.3958 | 1.9524 |
| 7 | 1.5332 | 2.1708 | 0.6376 | 1.8385 | 1.7450 | 2.1256 | 0.3805 | 1.9559 |
| 8 | 1.7654 | 2.5330 | 0.7676 | 2.1608 | 1.8027 | 2.1958 | 0.3931 | 1.9807 |
| 9 | 1.5877 | 2.3220 | 0.7343 | 1.9554 | 1.7619 | 2.1485 | 0.3866 | 1.9674 |
| 10 | 1.4749 | 2.2204 | 0.7455 | 1.8334 | 1.7586 | 2.1411 | 0.3825 | 1.9585 |
| 11 | 1.7139 | 2.5421 | 0.8282 | 2.1098 | 1.7973 | 2.1841 | 0.3868 | 1.9769 |
| 12 | 1.6753 | 2.4745 | 0.7993 | 2.0786 | 1.7926 | 2.1843 | 0.3917 | 1.9759 |
| 13 | 1.7974 | 2.6332 | 0.8357 | 2.2250 | 1.8014 | 2.2052 | 0.4039 | 1.9817 |
| 14 | 1.6354 | 2.4672 | 0.8318 | 2.0700 | 1.7925 | 2.1872 | 0.3947 | 1.9739 |
| 15 | 1.5482 | 2.1366 | 0.5884 | 1.8153 | 1.7536 | 2.1290 | 0.3754 | 1.9555 |

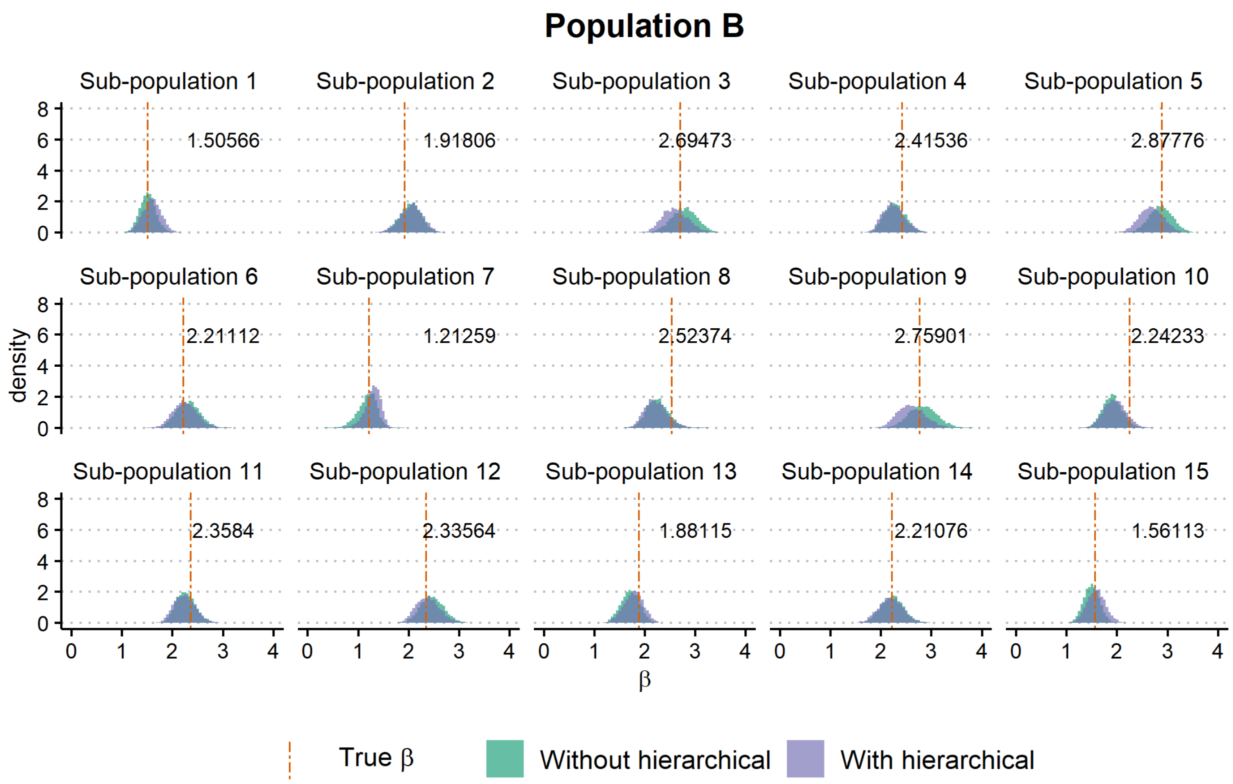


Figure S6: Marginal posteriors of $\beta$ for Population b

Table S2: Population B HPDs

| | Without a hierarchical model | | | | With a hierarchical model | | | |
|---|---|---|---|---|---|---|---|---|
| **Sub-population** | **HPD lower level** | **HPD upper level** | **Width of the HPD** | **Posterior median** | **HPD lower level** | **HPD upper level** | **Width of the HPD** | **Posterior median** |
| 1 | 1.2106 | 1.8168 | 0.6062 | 1.5094 | 1.2331 | 1.9254 | 0.6923 | 1.5907 |
| 2 | 1.6135 | 2.4081 | 0.7946 | 2.0450 | 1.6157 | 2.4875 | 0.8718 | 2.0571 |
| 3 | 2.2843 | 3.2463 | 0.9620 | 2.7915 | 2.1069 | 3.0890 | 0.9821 | 2.5947 |
| 4 | 1.8993 | 2.6731 | 0.7738 | 2.2713 | 1.8747 | 2.7014 | 0.8267 | 2.2424 |
| 5 | 2.4107 | 3.3125 | 0.9018 | 2.8650 | 2.2030 | 3.1286 | 0.9257 | 2.6731 |
| 6 | 1.8494 | 2.7329 | 0.8835 | 2.3067 | 1.8058 | 2.7179 | 0.9120 | 2.2433 |
| 7 | 0.7997 | 1.5139 | 0.7142 | 1.1990 | 0.9631 | 1.5650 | 0.6019 | 1.3041 |
| 8 | 1.8594 | 2.6858 | 0.8264 | 2.2429 | 1.8015 | 2.6622 | 0.8608 | 2.2068 |
| 9 | 2.3153 | 3.3710 | 1.0558 | 2.8279 | 2.0757 | 3.1230 | 1.0474 | 2.5921 |
| 10 | 1.5505 | 2.2991 | 0.7485 | 1.9050 | 1.5628 | 2.3903 | 0.8275 | 1.9502 |
| 11 | 1.8894 | 2.6358 | 0.7464 | 2.2525 | 1.8356 | 2.6521 | 0.8166 | 2.2338 |
| 12 | 2.0281 | 2.9156 | 0.8875 | 2.4603 | 1.9274 | 2.8457 | 0.9183 | 2.3754 |
| 13 | 1.3347 | 2.0570 | 0.7223 | 1.7166 | 1.3535 | 2.1228 | 0.7694 | 1.7837 |
| 14 | 1.7624 | 2.6569 | 0.8946 | 2.2216 | 1.7223 | 2.6568 | 0.9345 | 2.1957 |
| 15 | 1.1934 | 1.7933 | 0.5999 | 1.4974 | 1.2263 | 1.9511 | 0.7248 | 1.5828 |

##### S2.4 Testing for the presence of heterogeneity using the ROPE criterion

The Region of Practical Equivalence (ROPE) introduced by J. K. Kruschke (2013), coupled with a 95% HPD interval, can be used to formally test if a population is homogeneous or heterogeneous. A ROPE is a predefined range for the parameter values that cover most of the posterior distribution; " a small range of parameter values that are considered to be practically equivalent to the null value for purposes of the particular application" (J. Kruschke, 2014). A recent application of the ROPE criteria can be found in Shi et al. (2019) to test for invariance in a Bayesian factor analysis setting.

As a reference value for comparison with the posteriors, we took 2 (this is not because the true values of $\psi_\beta$ were equal to 2, but because the posterior medians of both populations' $\psi_\beta$ were close to 2). Then we used a common ROPE $(1.5, 2.5)$ for both populations to make a decision about the heterogeneity/ variability of the model parameter $\beta$ across the sub-populations in the populations. This step was implemented in R using the readily available package *BEST* (J. K. Kruschke & Meredith, 2020). Figures S7 and S8 illustrates the posteriors of $\beta$s of the sup-populations for the two populations with the ROPE criterion. For Population A, the pre-defined ROPE value completely included the 95% HPD intervals of all the sub-populations and this gives us enough evidence to conclude that the Population A has homogeneous sub-populations because 95% of most credible values of the sub-population specific $\beta$s are practically equivalent to the ROPE value we used. On the other hand, when we used the same ROPE value for Population B, the pre-defined ROPE value only partially included the 95% HPD intervals of all the sub-populations. This gives us enough evidence to conclude that the Population B has heterogeneous sub-populations because 95% of most credible values of the sub-population specific $\beta$s are variant with respect to the ROPE value we used. As Shi et al. (2019) mention, it must also be noted that choosing a ROPE is somewhat subjective but an unavoidable factor and the results (whether a population is homogeneous or heterogeneous) might be directly related to the value of the ROPE.

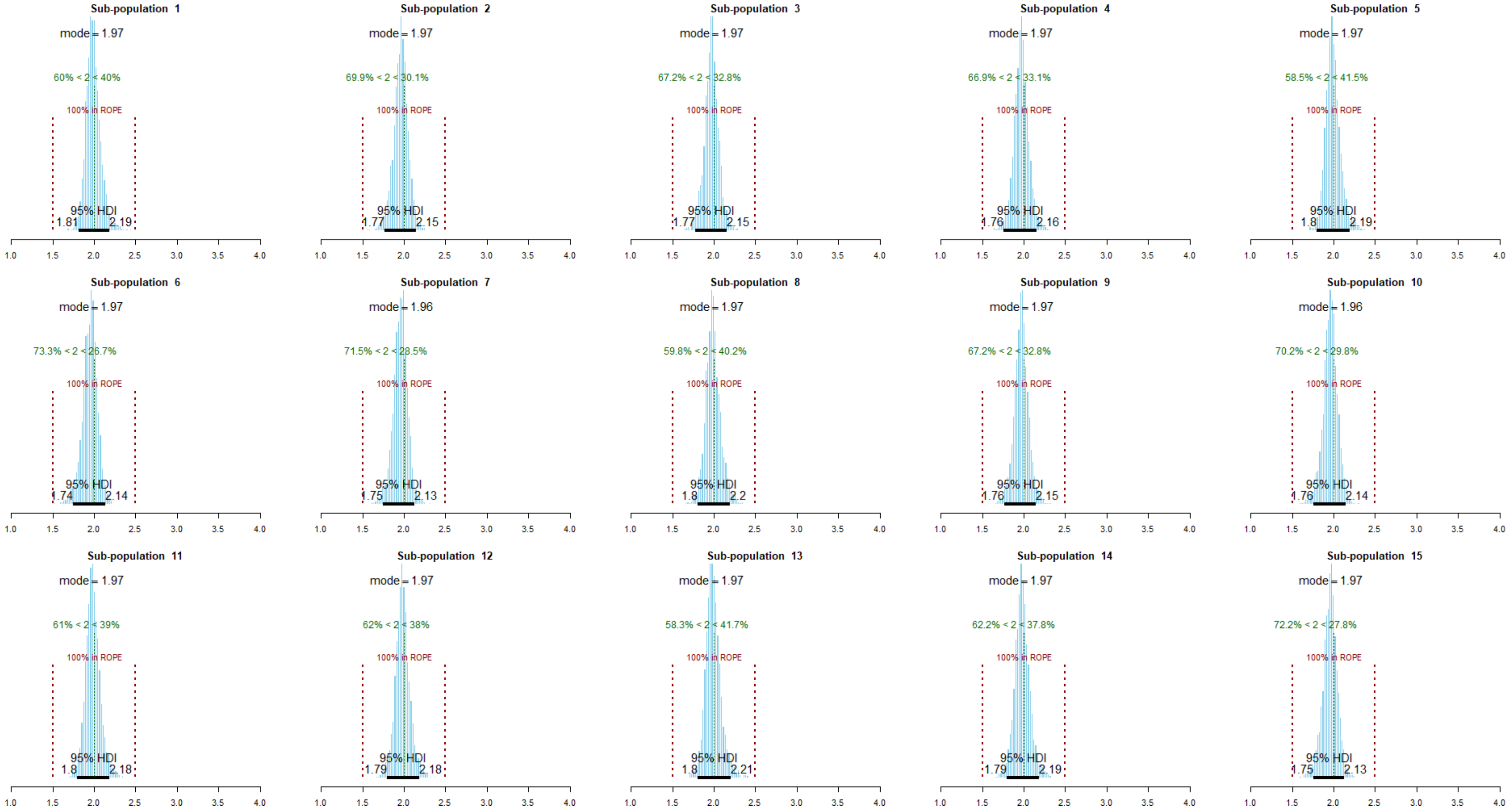


Figure S7: Marginal posteriors of $\beta_k$ (for $k = 1, 2, \ldots, 15$) for Population A with ROPE criteria. Here, 95%HDI (in black) refers to the 95% Highest Posterior Density Interval. The reference value for comparisons (i.e., 2) is shown as a green dashed line. The percentage of which the ROPE includes the HPD interval is shown in dark red. The posterior mode of each sub-population is also shown.

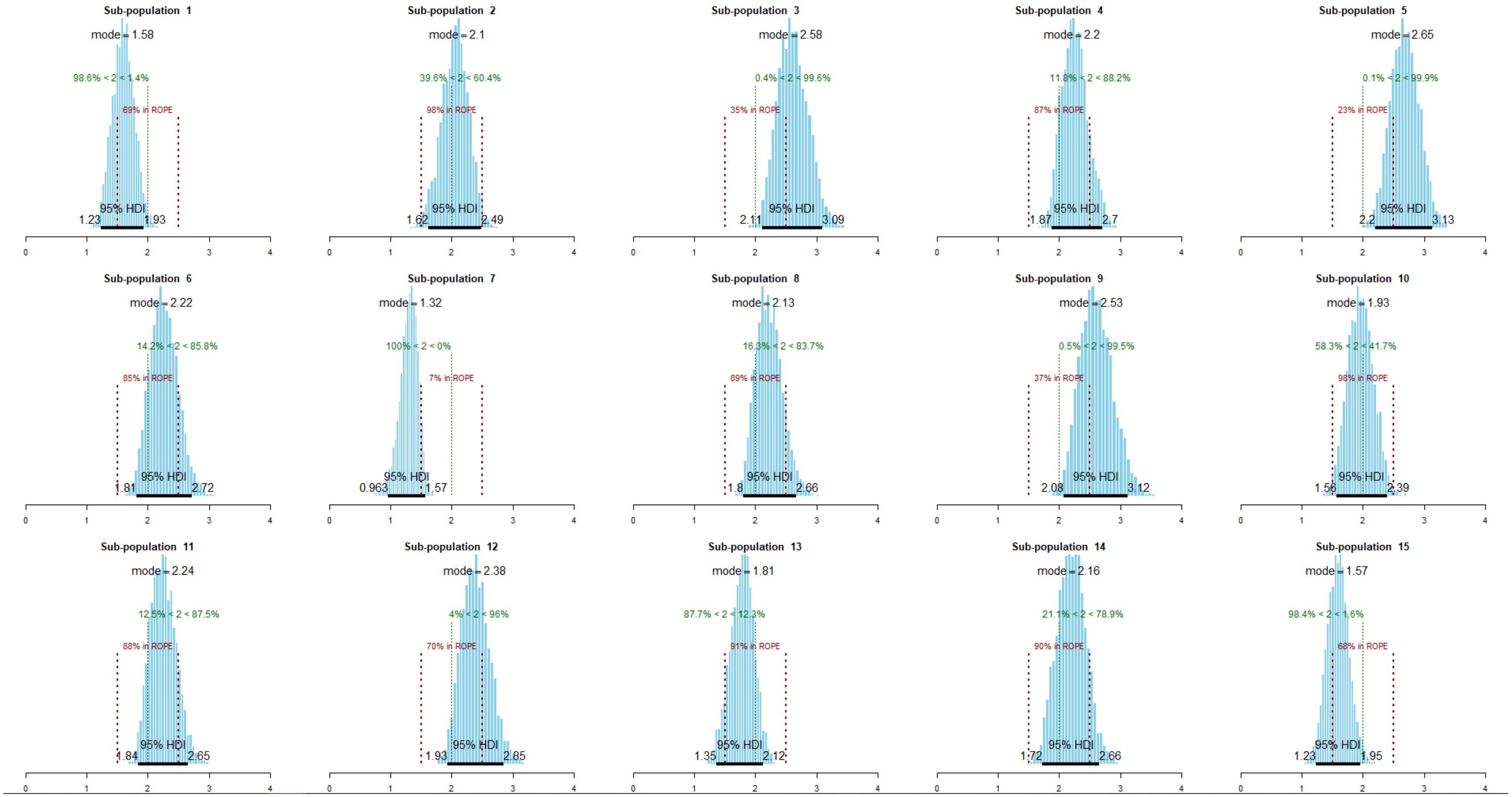


Figure S8: Marginal posteriors of $\beta_k$ (for $k = 1, 2, \ldots, 15$) for Population B with ROPE criteria. Here, 95%HDI (in black) refers to the 95% Highest Posterior Density Interval. The reference value for comparisons (i.e., 2) is shown as a green dashed line. The percentage of which the ROPE includes the HPD interval is shown in dark red. The posterior mode of each sub-population is also shown.

#### S3 Epidemic fade-out

##### S3.1 Criteria for observing a major outbreak

We first focused on an SIRS model with following model parameters and conditions:

1. $\beta = 2$ (rate of transmission).
2. $\gamma = 1$ (recovery rate).
3. $\mu = 0.06$ (rate at which the recovered individuals become susceptible again).
4. The sub-population size ($N$) is 1000.
5. Initially the epidemic starts with one infectious individual.

We generated a deterministic time plot of this SIRS model and identified a rough time interval where the first trough would be. Figure S9 illustrates this. The red dashed horizontal line is the endemic line for infectious individuals (28.3019) and points A $(12.44, 28.3019)$ and B $(29.01, 28.3019)$ are where the endemic line for infectious individuals and the deterministic path intersect. The time interval between A and B lines, denoted by blue dashed vertical lines, was considered to be the first trough. Therefore, we assumed that *if* we observe extinction in the first trough, we should observe it between A and B points. However, to allow for stochastic effects as well as the change of the epidemic dynamics due to the variability of the sub-population specific parameters, $\beta_k$ (for $k = 1, 2, \ldots, K$), we roughly defined that the stochastic system will enter the first trough —there is a major outbreak without an initial fade-out —when the number of infectious individuals at the 10th week or a later week(s) is either zero or has a decreasing trend in the number of infectious individuals than in the previous weeks.

After reaching the first trough, if the stochastic system did not reach zero infectious individuals by the time it reached 30 weeks and by that time, if the number of infectious individuals were still greater than 1, we considered that the stochastic system had exited the first trough (causing a second a second wave). If the system reaches zero infectious individuals by the time it reached 30 weeks or less, given that the system reached the first trough, we considered that the stochastic system will have an epidemic fade-out.

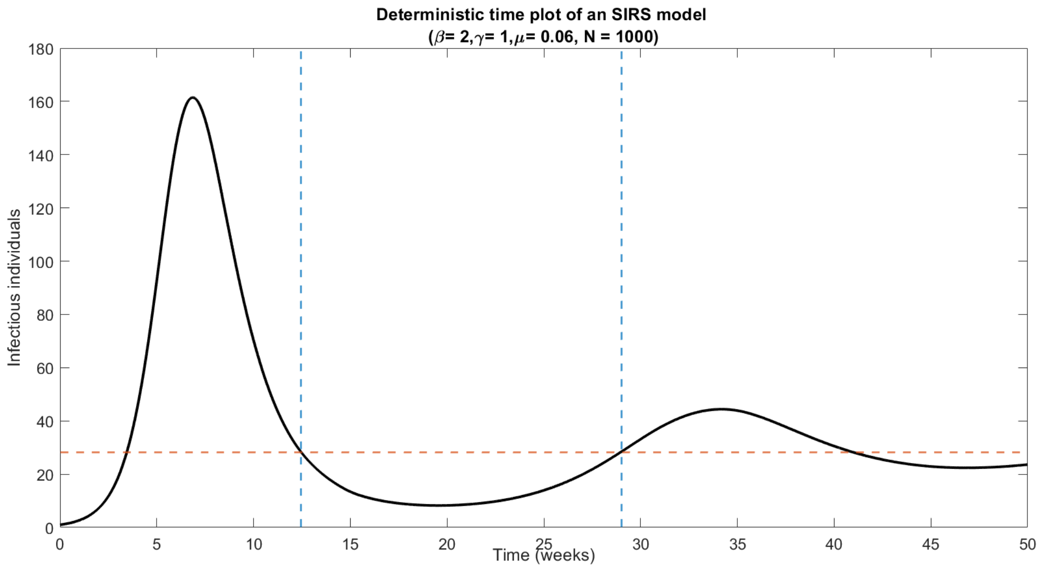


Figure S9: Deterministic time plot of an SIRS model with $(\beta, \gamma, \mu = 2, 1, 0.06)$ for sub-population of size 1000 and initially, one infectious individual.

##### S3.2 Estimating the probability of epidemic fade-out

We used methods introduced by Ballard, Bean, and Ross (2016) at this stage. However, to avoid the repetition of calculating the probabilities given similar transmission rate ranges, we first considered transmission rates values in the interval $(1.07, 4)$ and calculated the corresponding probabilities of epidemic fade-out for an $SIRS$ model with these values. When specific regions needed attention, we used the MATLAB function *interp1* to interpolate in the required transmission rate region; the sampled posterior values. Furthermore, when plots illustrating the relationship between transmission rate and the probability of epidemic fade-out are plotted before plotting all the probabilities were interpolated across 5000 points within $(1.07, 5)$ transmission rate range. The starting point as 1.07 was used since the probability of epidemic fade-out is incalculable using methods by Ballard et al. (2016) when the transmission rate dropped below 1.07. Hence, the sampled values for $\beta < 1.07$ from the posteriors for $\beta_k$ were ignored when calculating the epidemic fade-out probabilities.

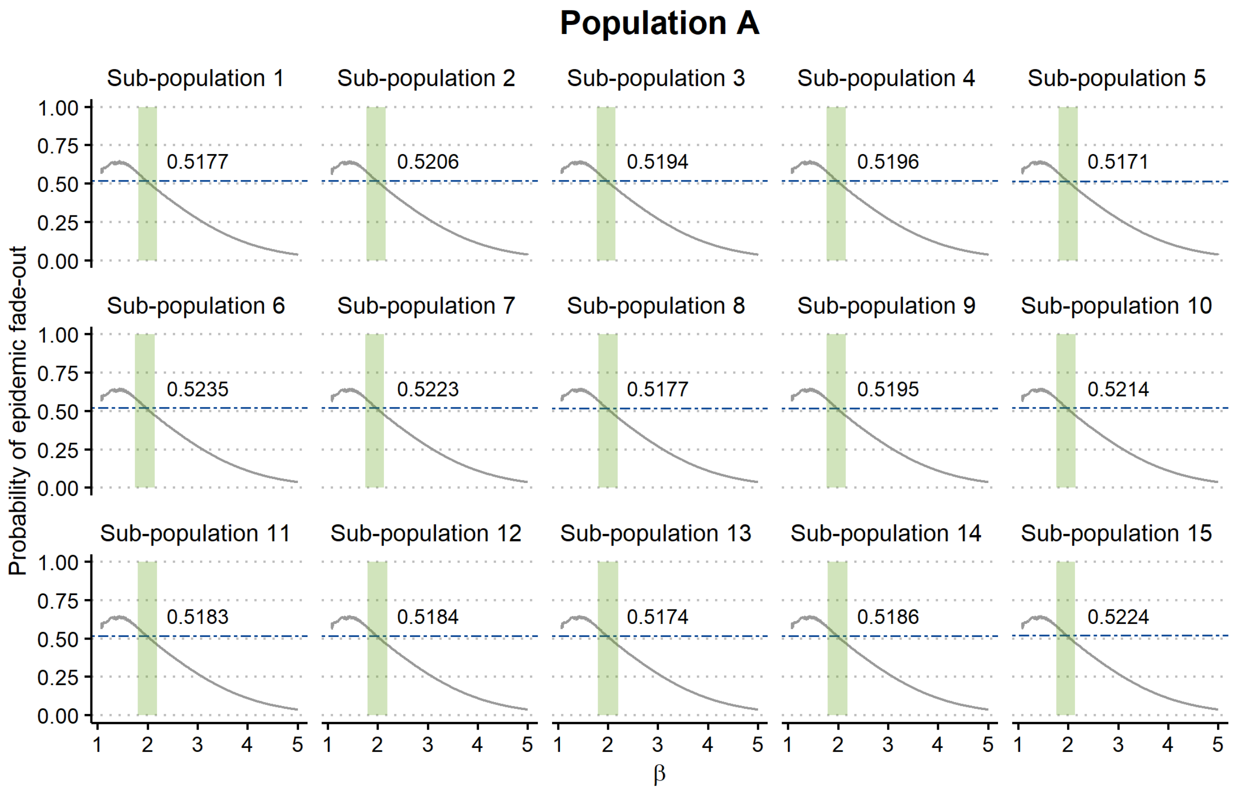


Figure S10: **Population A:** Change of probability of epidemic fade-out with respect to $\beta$ (in grey). The green solid areas are the 95% HPD intervals of the posterior of $\beta_k$ under each sub-population. The blue dashed lines are the medians and their values are the median probabilities of epidemic fade-out.

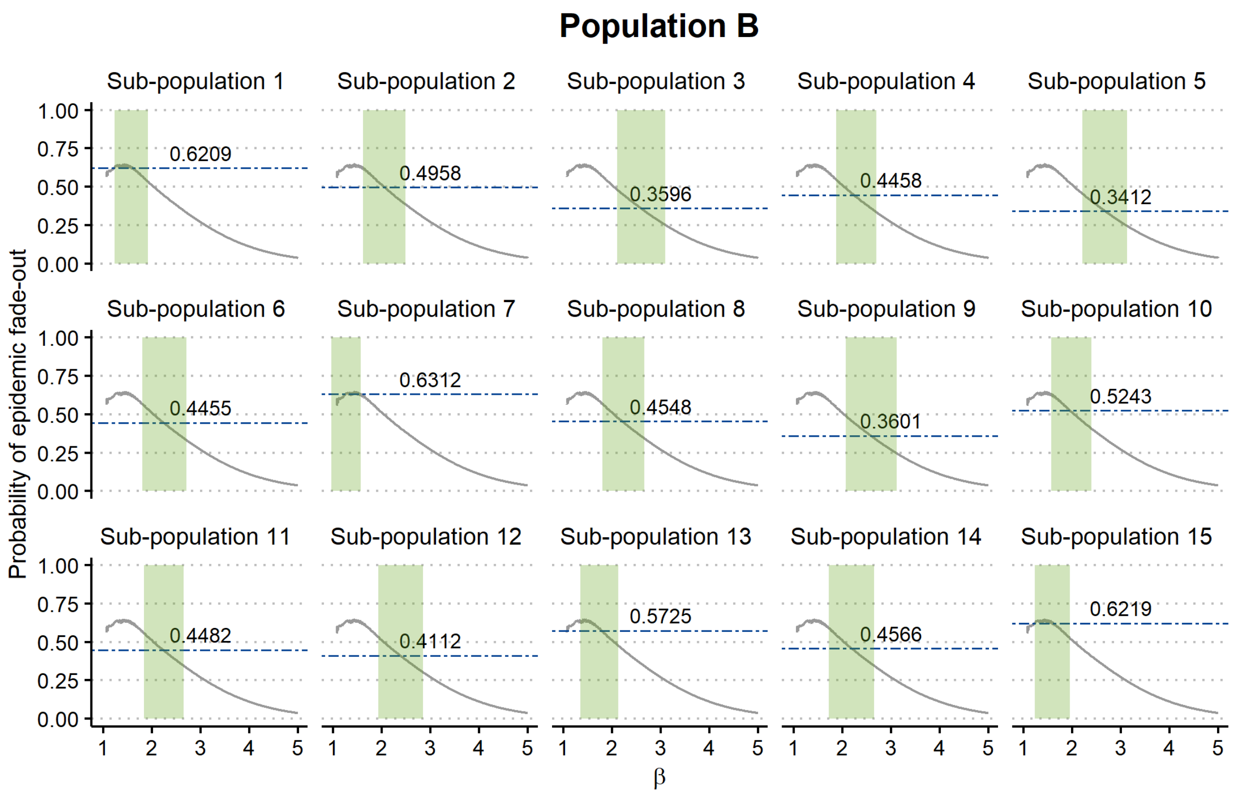


Figure S11: **Population B:** Change of probability of epidemic fade-out with respect to $\beta$ (in grey). The green solid areas are the 95% HPD intervals of the posterior of $\beta_k$ under each sub-population. The blue dashed lines are the medians and their values are the median probabilities of epidemic fade-out.

### S4 Accuracy of estimates under a hierarchical modelling framework

#### S4.1 Marginal posterior distributions of the hyper-parameters of 16 sub-populations (using the two-step methodology)

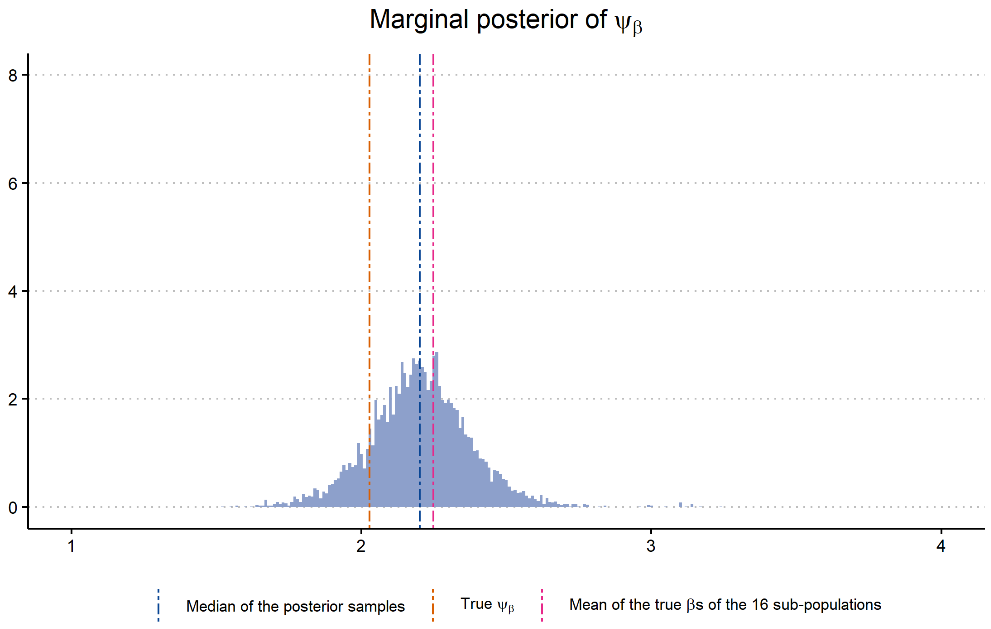


Figure S12: Marginal posterior of $\psi_\beta$ 16 sub-populations .

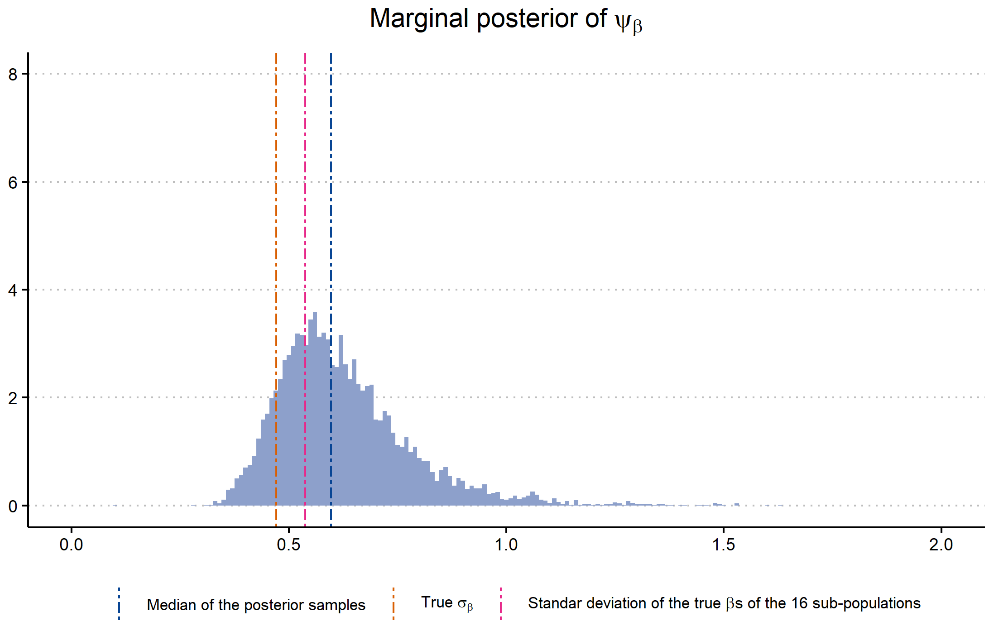


Figure S13: Marginal posterior of $\sigma_\beta$ 16 sub-populations

##### S4.2 Comparison of sub-populations specific parameters under a hierarchical model (Step 2 of the two-step methodology) and independently estimated values.

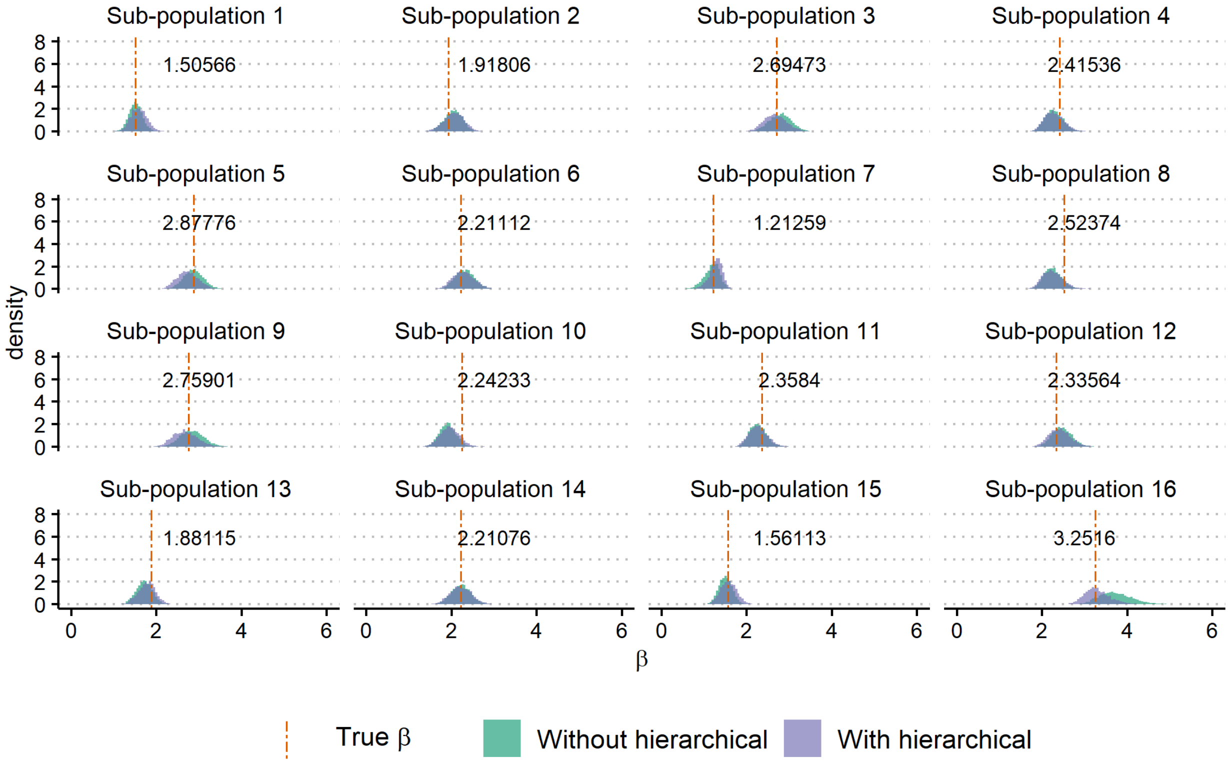


Figure S14: Marginal posteriors of $\beta$ 16 sub-populations

Table S3: Population C HPDs

| | Without a hierarchical model | | | | With a hierarchical model | | | |
|---|---|---|---|---|---|---|---|---|
| Sub-population | HPD lower level | HPD upper level | Width of the HPD | Posterior median | HPD lower level | HPD upper level | Width of the HPD | Posterior median |
| 1 | 1.2106 | 1.8168 | 0.6062 | 1.5094 | 1.2204 | 1.9474 | 0.7270 | 1.5821 |
| 2 | 1.6135 | 2.4081 | 0.7946 | 2.0450 | 1.5929 | 2.4683 | 0.8755 | 2.0618 |
| 3 | 2.2843 | 3.2463 | 0.9620 | 2.7915 | 2.1621 | 3.1590 | 0.9969 | 2.6630 |
| 4 | 1.8993 | 2.6731 | 0.7738 | 2.2713 | 1.8582 | 2.7133 | 0.8551 | 2.2695 |
| 5 | 2.4107 | 3.3125 | 0.9018 | 2.8650 | 2.2545 | 3.2278 | 0.9733 | 2.7294 |
| 6 | 1.8494 | 2.7329 | 0.8835 | 2.3067 | 1.8084 | 2.7551 | 0.9466 | 2.2758 |
| 7 | 0.7997 | 1.5139 | 0.7142 | 1.1990 | 0.9301 | 1.5442 | 0.6141 | 1.2856 |
| 8 | 1.8594 | 2.6858 | 0.8264 | 2.2429 | 1.8379 | 2.7399 | 0.9020 | 2.2385 |
| 9 | 2.3153 | 3.3710 | 1.0558 | 2.8279 | 2.1390 | 3.2278 | 1.0888 | 2.6620 |
| 10 | 1.5505 | 2.2991 | 0.7485 | 1.9050 | 1.5412 | 2.3851 | 0.8439 | 1.9577 |
| 11 | 1.8894 | 2.6358 | 0.7464 | 2.2525 | 1.8738 | 2.7238 | 0.8499 | 2.2561 |
| 12 | 2.0281 | 2.9156 | 0.8875 | 2.4603 | 1.9600 | 2.9317 | 0.9717 | 2.4051 |
| 13 | 1.3347 | 2.0570 | 0.7223 | 1.7166 | 1.3502 | 2.1396 | 0.7894 | 1.7748 |
| 14 | 1.7624 | 2.6569 | 0.8946 | 2.2216 | 1.7388 | 2.7089 | 0.9701 | 2.2199 |
| 15 | 1.1934 | 1.7933 | 0.5999 | 1.4974 | 1.2041 | 1.9175 | 0.7135 | 1.5745 |
| 16 | 3.0260 | 4.5087 | 1.4827 | 3.7093 | 2.7450 | 3.9093 | 1.1643 | 3.2642 |

#### S5 Additional figures and details

##### S5.1 Spread of the true $\beta$ values within the Truncated Normal$(2, 0.5^2, 1, 10)$

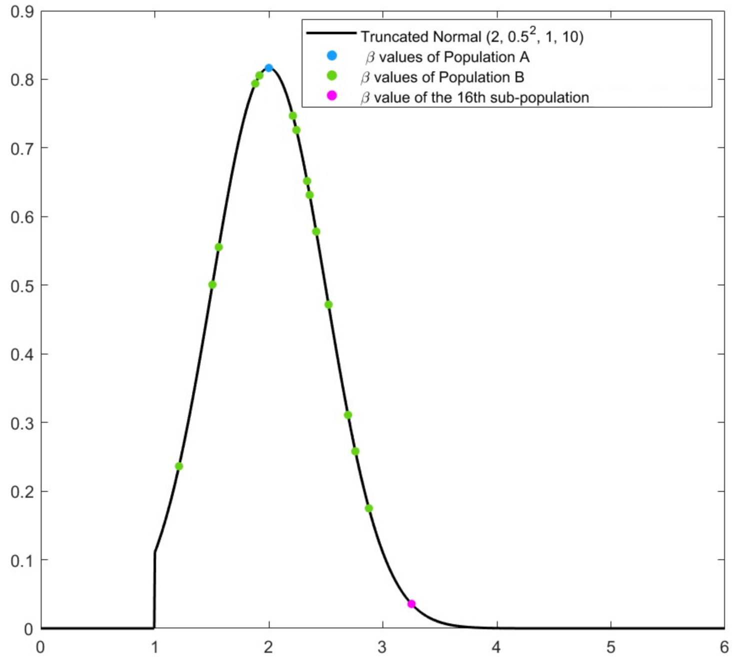


Figure S15: Truncated Normal$(2, 0.5^2, 1, 10)$ in black. Blue dot represents the true $\beta = 2$ of all the sub-population in Population A. Green dots represents the $\beta_k$ values in the sub-populations in Population B and the first 15 sub-populations. The magenta dot represents the true $\beta_{16}]$ .

##### S5.2 MATLAB Codes

The codes can be found on GitHub at:
`https://github.com/PunyaAlahakoon/Hierarchical_parameter_estimation.git`
